# Utility of polygenic embryo screening for disease depends on the selection strategy

**DOI:** 10.1101/2020.11.05.370478

**Authors:** Todd Lencz, Daniel Backenroth, Einat Granot-Hershkovitz, Adam Green, Kyle Gettler, Judy H. Cho, Omer Weissbrod, Or Zuk, Shai Carmi

## Abstract

Polygenic risk scores (PRSs) have been offered since 2019 to screen *in vitro* fertilization embryos for genetic liability to adult diseases, despite a lack of comprehensive modeling of expected outcomes. Here we predict, based on the liability threshold model, the expected reduction in complex disease risk following *polygenic embryo screening* for a single disease. Our main finding is that a strong determinant of the potential utility of such screening is the *selection strategy*, a factor that has not been previously studied. Specifically, when only embryos with a very high PRS are excluded, the achieved risk reduction is minimal. In contrast, selecting the embryo with the lowest PRS can lead to substantial relative risk reductions, given a sufficient number of viable embryos. For example, a relative risk reduction of ≈50% for schizophrenia can be achieved by selecting the embryo with the lowest PRS out of five viable embryos. We systematically examine the impact of several factors on the utility of screening, including the variance explained by the PRS, the number of embryos, the disease prevalence, the parental PRSs, and the parental disease status. When quantifying the utility, we consider both relative and absolute risk reductions, as well as population-averaged and per-couple risk reductions. We also examine the risk of pleiotropic effects. Finally, we confirm our theoretical predictions by simulating “virtual” couples and offspring based on real genomes from schizophrenia and Crohn’s disease case-control studies. We discuss the assumptions and limitations of our model, as well as the potential emerging ethical concerns.

## Introduction

Polygenic risk scores (**PRS**s) have become increasingly well-powered, relying on findings from large-scale genome-wide association studies for numerous diseases (Visscher et al., 2017; Wray et al., 2013). Consequently, a growing body of research has examined the potential clinical utility of applying PRSs in the treatment of adult patients in order to identify those at heightened risk for common late-onset diseases such as coronary artery disease or breast cancer (Britt et al., 2020; Khera et al., 2018; Torkamani et al., 2018). Another potential application of PRSs is preimplantation screening of *in vitro* fertilization (**IVF**) embryos, or *polygenic embryo screening* (**PES**). Polygenic embryo screening has been offered since 2019 (Treff, Eccles, et al., 2019), but has been the focus of comparatively little empirical research, despite debate over ethical and social concerns surrounding the practice (Anomaly, 2020; Lázaro-Muñoz et al., 2020; Munday & Savulescu, 2021).

We have recently demonstrated that screening embryos on the basis of polygenic scores for quantitative traits (such as height or intelligence) has limited utility in most realistic scenarios (Karavani et al., 2019), and that the accuracy of the score is a more significant determinant of PES utility for quantitative traits compared with the number of available embryos. On the other hand, a series of four studies (Lello et al., 2020; Treff, Eccles, et al., 2019; Treff et al., 2020; Treff, Zimmerman, et al., 2019) conducted by a private company providing PES services has suggested that PES for dichotomous disease risk may have significant clinical utility. However, these studies examined a relatively limited range of scenarios, primarily focusing on distinctions between sibling pairs discordant for illness, and did not provide a comprehensive examination of various potential PES settings. Filling this gap is an urgent need, as understanding the statistical properties of PES forms a critical foundation to any ethical consideration (Lázaro-Muñoz et al., 2020).

Here, we use statistical modeling to examine the potential utility of PES for reducing disease risk, with an aim toward informing future ethical deliberations. We focus on screening for a single complex disease, and study a range of realistic scenarios, quantifying the role of parameters such as the variance explained by the score, the number of available embryos, and the disease prevalence. We show that a major determinant of the outcome of PES is the *selection strategy*, namely the way in which an embryo is selected for implantation given the distribution of PRSs across embryos. We also study the risk reduction *conditional* on parental PRSs or disease status, and consider the risk of developing diseases not screened. Finally, we validate some of our predictions based on real genomes of cases and controls for two common complex diseases.

## Results

### Model and selection strategies

For each analysis presented below, we assume that a couple has generated, by IVF, *n* viable embryos such that each embryo, if implanted, would have led to a live birth. We focus on a single complex disease, and assume that the corresponding PRS has been computed for each embryo. Given the PRSs of the *n* embryos, a single embryo is selected for implantation based on a *selection strategy*.

The first strategy we consider is aimed only at avoiding high-risk embryos, consistent with studies of the potential clinical utility of PRSs in adults (Chatterjee et al., 2016; Dai et al., 2019; Gibson, 2019; Khera et al., 2018; Mars et al., 2020; Mavaddat et al., 2019; Torkamani et al., 2018). For example, the first case report presented on PES described the identification and exclusion of embryos with extremely high (top 2-percentiles) PRS (Treff, Eccles, et al., 2019). We term this strategy “*high-risk exclusion*” (**HRE**: **Figure 1A**, upper panel). Under HRE, after high-risk embryos are set aside, an embryo is randomly selected for implantation among the remaining available embryos. (In the case that all embryos are high-risk, we assume a random embryo is selected among them.)

**Figure 1.**
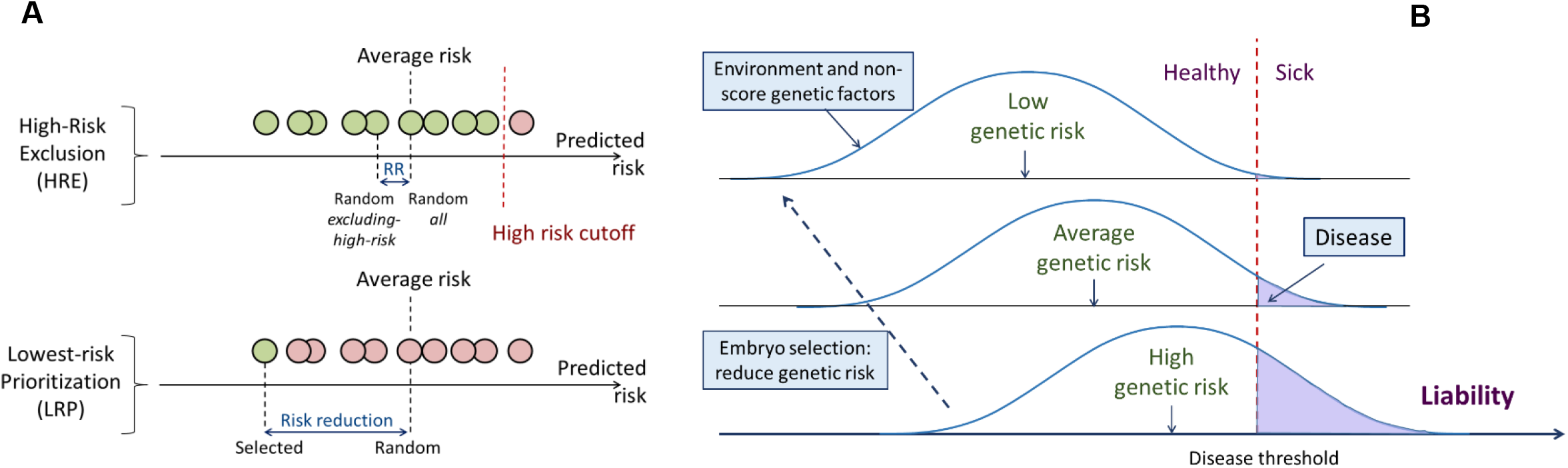
A schematic of the liability threshold model and polygenic embryo screening. (A) An illustration of the embryo selection strategies considered in this report. In the figure, each embryo is shown as a filled circle, and embryos are sorted based on their predicted risk, i.e., their polygenic risk scores. Excluded embryos are shown in pink, and embryos that can be implanted in green. The risk reduction (RR) is indicated as the difference in risk between a randomly selected embryo (if no polygenic scoring was performed) and the embryo selected based on one of two strategies. In *high-risk exclusion* (HRE), the embryo selected for implantation is random, as long as its PRS is under a high-risk cutoff (usually the top few PRS percentiles). If all embryos are high-risk, a random embryo is selected. In *lowest-risk prioritization* (LRP), the embryo with the lowest PRS is selected for implantation. As we describe below, the LRP strategy yields much larger disease risk reductions. (B) An illustration of the liability threshold model (LTM). Under the LTM, each disease has an underlying (unobserved) liability, and an individual is affected if the total liability is above a threshold. The liability is composed of a genetic component and an environmental component, both assumed to be normally distributed in the population. For a given genetic risk (represented here by the polygenic risk score), the liability is the sum of that risk, plus a normally distributed *residual* component (environmental + genetic factors not captured by the PRS). For an individual with high genetic risk (bottom curve), even a modestly elevated (and thus, commonly-occurring) liability-increasing environment will lead to disease. For an individual with low genetic risk (top curve), only an extreme environment will push the liability beyond the disease threshold. Thus, disease risk reduction can be achieved with embryo screening by lowering the genetic risk of the implanted embryo. (Note that for the purpose of illustration, panel (B) displays three discrete levels of genetic risk, although in reality, the PRS is continuously distributed.)

An alternative selection strategy is to use the embryo with the lowest PRS. Ranking and prioritizing embryos for implantation based on morphology is common in current IVF practice (Bormann et al., 2020; Montag et al., 2013; Rhenman et al., 2015). If ranking is instead based on a disease PRS, the embryo with the lowest PRS could be selected, without any recourse to high-risk PRS thresholds. Such an approach was suggested by another recent publication from the same company (based on a multi-disease index), but statistical comparisons were only examined in the context of sibling pairs (Treff et al., 2020). We term the implantation of the embryo with the lowest PRS as “*lowest-risk-prioritization*” (**LRP**; **Figure 1A**, lower panel).

In the following, we describe the theoretical risk reduction that can be achieved under these selection strategies. Our statistical approach is based on the liability threshold model (**LTM**; (Falconer, 1967)). The LTM represents disease risk as a continuous liability, comprising genetic and environmental risk factors, under the assumption that individuals with liability exceeding a threshold are affected. The liability threshold model has been shown to be consistent with data from family-based transmission studies (Wray & Goddard, 2010) and GWAS data (Visscher & Wray, 2015). Consequently, we define the disease risk of a given embryo probabilistically, as the chance that, given its PRS, its liability will cross the threshold at any point after birth (**Figure 1B**).

We use the following notation. We define the predictive power of a PRS as the proportion of variance in the liability of the disease explained by the score (Dudbridge, 2013), and denote it as 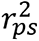. We quantify the outcome of PES in two ways: the *relative risk reduction* (**RRR**) is defined as 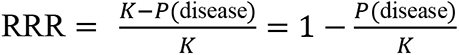, where *K* is the disease prevalence and *P*(*disease*) is the probability of the selected embryo to be affected; the *absolute risk reduction* (**ARR**) is defined as *K* − *P*(*disease*). For example, if a disease has prevalence of 5% and the selected embryo has a probability of 3% to be affected, the RRR is 40%, and the ARR is 2% points. We computed the RRR and ARR analytically under each selection strategy, and for various values for the disease prevalence, the strength of the PRS, embryo exclusion thresholds, and other parameters. The mathematical details of the calculations are provided in the *Materials and Methods*.

### The risk reduction under the *high-risk exclusion* strategy

In **Figure 2** (upper row), we show the relative risk reduction achievable under the HRE strategy with *n* = 5 embryos. Under the 2-percentile threshold (straight black lines), the reduction in risk is limited: the RRR is <10% in all scenarios where 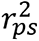 ≤ 0.1. Currently, 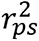 ≈ 0.1 (on the liability scale) is the upper limit of the predictive power of PRSs for most complex diseases (Lambert et al., 2021), with the exception of a few disorders with large-effect common variants (such as Alzheimer’s disease or type 1 diabetes) (Sharp et al., 2019; Q. Zhang et al., 2020). In the future, more accurate PRSs are expected. However, the common-variant SNP heritability is at most ≈30% even for the most heritable diseases such as schizophrenia and celiac disease (Holland et al., 2020; Y. Zhang et al., 2018), and it was recently suggested that 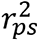 = 0.3 is the maximal realistic value for the foreseeable future (Wray et al., 2020). At this value, relative risk reduction would be 20% for *K* = 0.01, 9% for *K* = 0.05, and 3% for *K* = 0.2. These gains achieved with HRE are small because the overwhelming majority of affected individuals do not have extreme scores (Murray et al., 2020; Wald & Old, 2019).

**Figure 2.**
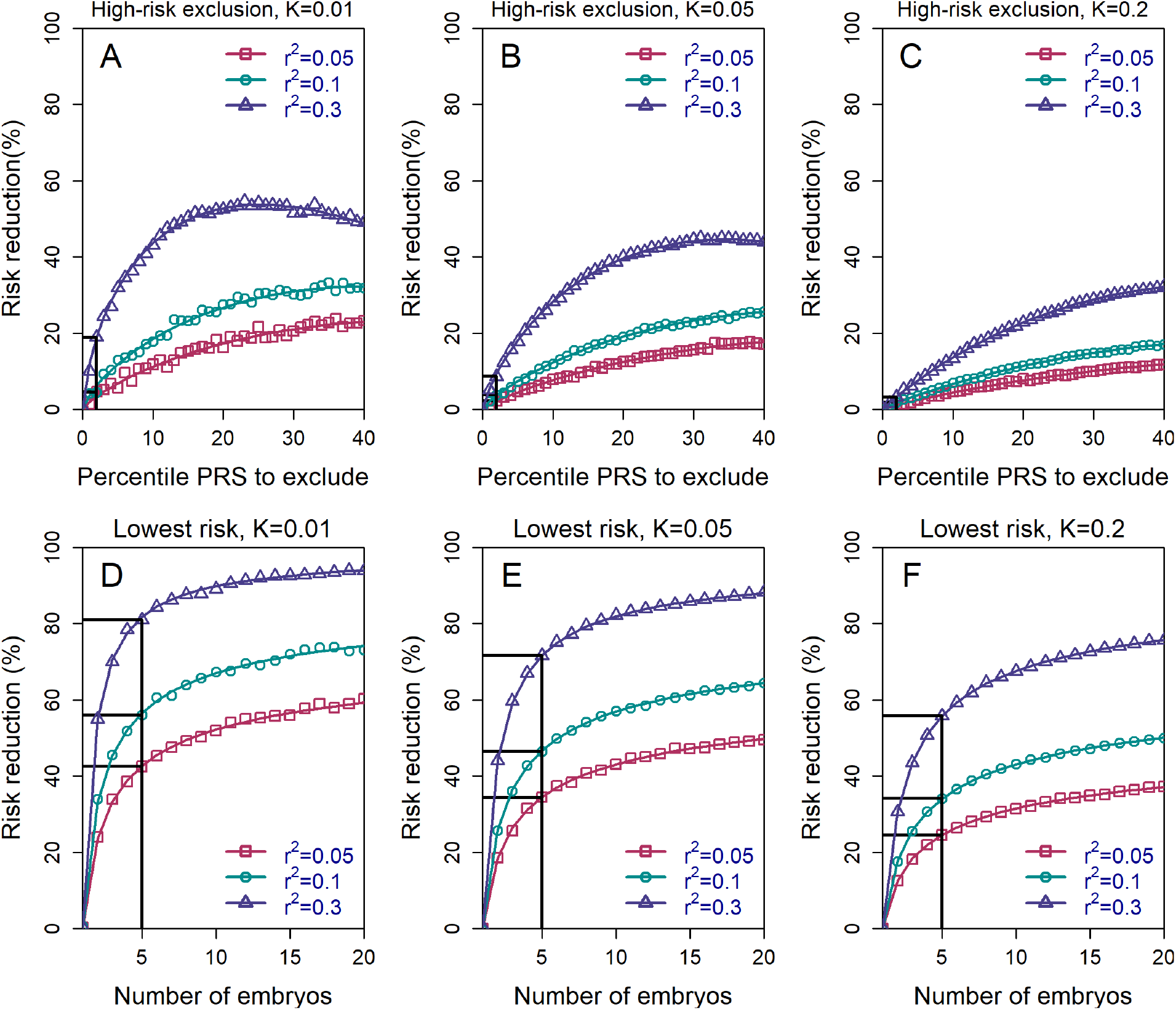
The relative risk reduction across selection strategies and disease parameters. The relative risk reduction (RRR) is defined as (*K* − *P*(disease))/*K*, where *K* is the disease prevalence, and *P*(disease) is the probability of the implanted embryo to become affected. The RRR is shown for the *high-risk exclusion* (HRE) strategy in the upper row (panels (A)-(C)), and for the *lowest-risk prioritization* (LRP) in the lower row (panels (D)-(F)). See Figure 1 for the definitions of the strategies. Results are shown for values of *K* = 0.01, 0.05, and 0.2 (panels (A)-(C), respectively), and within each panel, for variance explained by the PRS (on the liability scale) 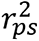 = 0.05, 0.1, and 0.3 (legends). Symbols denote the results of Monte-Carlo simulations (*Materials and Methods*), where PRSs of embryos were drawn based on a multivariate normal distribution, assuming PRSs are standardized to have zero mean and variance 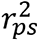, and accounting for the genetic similarity between siblings (Eq. (4) in *Materials and Methods*). In each simulated set of *n* sibling embryos (*n* = 5 for all simulations under HRE), one embryo was selected according to the selection strategy. The liability of the selected embryo was computed by adding a residual component (drawn from a normal distribution with zero mean and variance 1 − 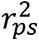) to its polygenic score. The embryo was considered affected if its liability exceeded *z*_*K*_, the (upper) *K*-quantile of the standard normal distribution. We repeated the simulations over 10^6^ sets of embryos and computed the disease risk. In each panel, curves correspond to theory: Eq. (31) in *Materials and Methods* for the HRE strategy, and Eq. (20) in *Materials and Methods* for the LRP strategy. Black straight lines correspond to the RRR achieved when excluding embryos at the top 2% of the PRS (for HRE, upper panels) or for selecting the lowest risk embryo out of *n* = 5 (for LRP, lower panels).

Risk reduction increases as the threshold for exclusion is expanded to include the top quartile of scores, and then reaches a maximum at ≈25-50% under a range of prevalence and 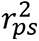 values. For all of these simulations, we set the number of available (testable) embryos to *n* = 5 (Dahdouh, 2021; Sunkara et al., 2011), although we acknowledge that the number of viable embryos may be much lower for many couples seeking IVF services for infertility (Smith et al., 2015). Simulations show that these estimates do not change much with increasing the number of embryos (see **Figure 2 - Figure Supplement 1**). This holds especially at more extreme threshold values, since most batches of *n* embryos will not contain any embryos with a PRS within, e.g., the top 2-percentiles.

It should be noted that the relative risk reduction does not increase monotonically under HRE. Under our definition, whenever all embryos are high risk, an embryo is selected at random. Thus, at the extreme case when all embryos (i.e, top 100%) are designated as high risk, an embryo is selected at random at all times, and the relative risk reduction reduces to zero. We chose this definition of the HRE strategy because it does not involve ranking of the embryos. However, we can also consider an alternative strategy: if all embryos are high risk, the embryo with the lowest PRS is selected. Here, the RRR is expected to increase when increasing the threshold and designating more embryos as high risk, which we confirm in **Figure 2 – Figure Supplement 2**. When the threshold is at 100% (all embryos are high risk), this alternative strategy (which we do not further consider) reduces to the *low-risk prioritization* strategy, which we study next.

### The risk reduction under the *low-risk prioritization* strategy

The HRE strategy treats all *non*-high-risk embryos equally. In practice, we expect most, or even all, embryos to be designated as non-high-risk, given the recent focus on the top PRS percentiles in the literature (e.g., Khera et al. 2018). However, as we have seen, this strategy leads to very little risk reduction. In **Figure 2** (lower panels), we show the expected RRR for the *low-risk prioritization* strategy, under which we prioritize for implantation the embryo with the lowest PRS, regardless of any PRS cutoff. Indeed, under the LRP strategy, risk reductions are substantially greater than in HRE. For example, with *n* = 5 available embryos, RRR>20% across the entire range of prevalence and 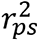 parameters considered, and can reach ≈50% for *K* ≤ 5% and 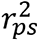 = 0.1, and even ≈80% for *K* = 1% and 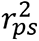 = 0.3. While RRR continues to increase as the number of available embryos increases, the gains are quickly diminishing after *n* = 5. On the other hand, Figure 2 also demonstrates that RRR drops steeply if the number of embryos falls below *n* = 5, although the lower bound for RRR when just two embryos are available (≈20% for many scenarios) is still comparable to the upper bound of the HRE strategy for a greater number of embryos.

### Effects of PES on dichotomous vs quantitative traits

Our results demonstrate that, contrary to our previous study reporting only small effects of PES for quantitative traits (Karavani et al., 2019), PES can generate substantial relative risk reductions for disease under the LRP strategy. To understand the relation between continuous and binary traits, consider an example involving IQ. Our estimate for the mean gain in IQ that could be achieved by selecting the embryo with the highest IQ polygenic score is approximately ≈2.5 IQ points (Karavani et al. 2019). Now assume that individuals with IQ<70 (2 SDs below the mean) are considered “affected” according to a dichotomized trait of “cognitive impairment.” Among individuals with IQ<70, the proportion of individuals with IQ in the range [67.5,70] is 33.5% (assuming a normal distribution). A gain of 2.5 points would shift such offspring beyond the threshold for “cognitive impairment,” resulting in a corresponding 33.5% reduction in risk of being “affected”. (Note that the above explanation is intended to provide an intuition and ignores any variability in the gain.) **Figure 2 – Figure Supplement 3** utilizes statistical modeling (with 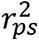 derived from recent GWAS for intelligence (Savage et al., 2018)) to demonstrate that substantial risk reductions can be achieved for a dichotomized trait, including when selecting out of just three embryos (panel (A)). Panel (B) extends these results to data for LDL cholesterol (with 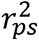 derived from (Weissbrod et al., 2021)); given *n* = 5 embryos and the currently available PRS for LDL-C levels, risk reductions for “high cholesterol” range from 40-60%, depending on the LDL level used to define the categorical trait. Thus, while implanting the embryo with the most favorable PRS is expected to result in very modest gains in an underlying quantitative trait, it is at the same time effective in avoiding embryos at the unfavorable tail of the trait.

### Effects of parental PRS and disease status

We next examined the effects of parental PRSs on the achievable risk reduction (*Materials and Methods*), given that families with high genetic risk for a given disease may be more likely to seek PES. **Figure 3** demonstrates that, as expected, the HRE strategy shows greater relative risk reduction as parental PRS increases, in particular when excluding only very high-scoring embryos. This result follows directly from the fact that, on average, offspring will tend to have PRS scores near the mid-parental PRS value. In contrast, the relative RR (though not the *absolute* RR; see next section) for the LRP strategy somewhat declines as parental PRSs increase. Nevertheless, the RRR for the LRP strategy remains greater than that for the HRE strategy across all parameters (as expected by the definitions of these strategies).

**Figure 3.**
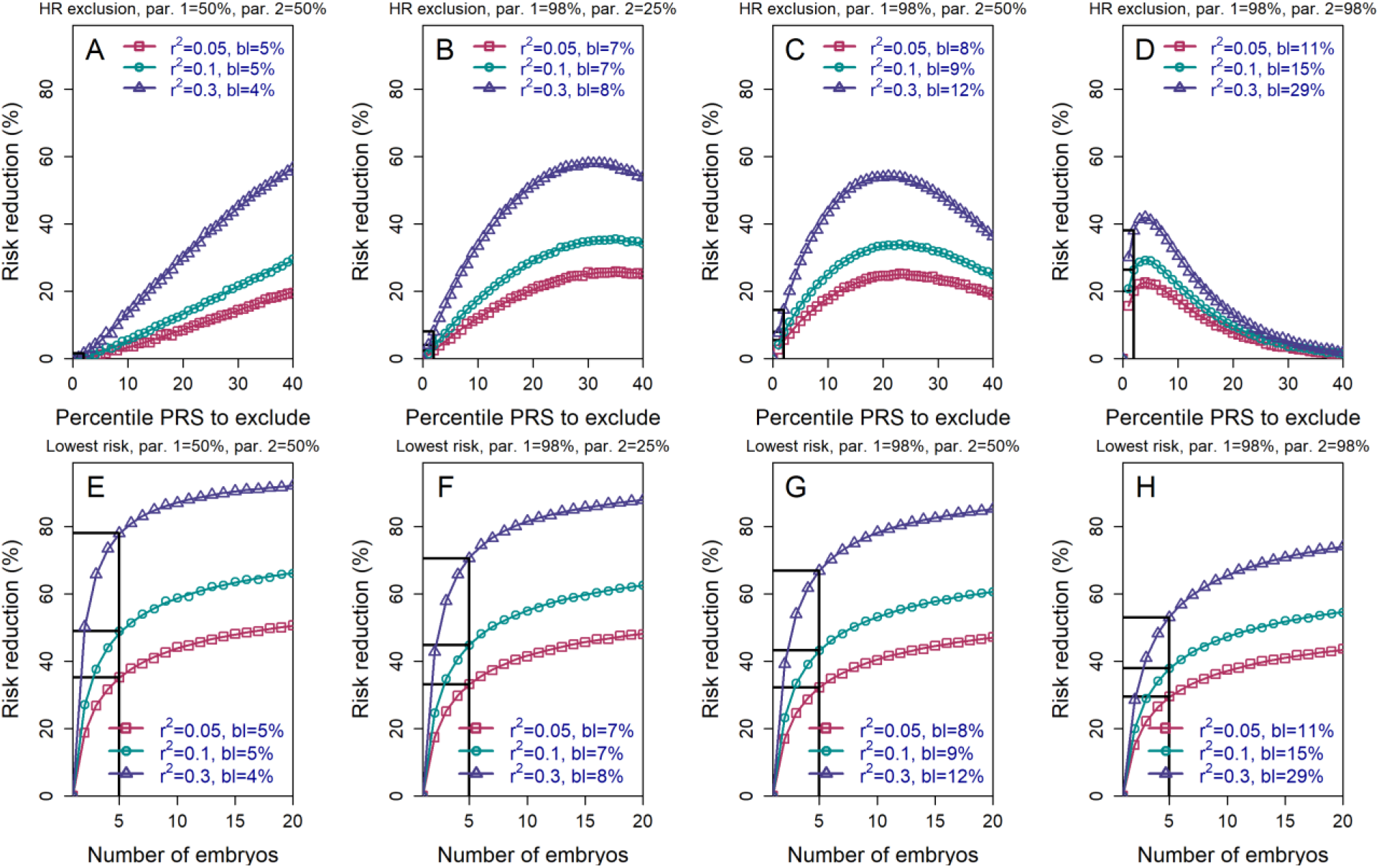
The relative risk reduction when the polygenic risk scores of the parents are known. Panels (A)-(D) are for the *high-risk exclusion* (HRE) strategy, while panels (E)-(H) are for the *lowest-risk prioritization* (LRP) strategy. All details are as in Figure 2, except the following. First, we fixed the prevalence to *K* = 5%. Second, in the simulations, we drew the PRS of each embryo as *s*_*i*_ = *x*_*i*_ + *c* (*i* = 1, …, *n*), where *x*_*i*_ is an embryo-specific component (independent across embryos) and *c* is the shared component, also representing the mean parental PRS (*Materials and Methods*). This is so far as in Figure 2; however, here we assumed that *c* is given, equal to the average PRSs of the two parents. In each panel, we consider a different pair of PRSs for the parents. For example, in panels (A) and (E), both parents (“par. 1” and “par. 2”) have PRS equal to the 50% percentile of the PRS distribution; in panels (B) and (F), one parent has PRS equal to the 98% percentile of the PRS distribution, while the other has PRS equal to the 25% percentile; and so on. Third, in the simulations, we computed the risk reduction (according to either strategy) relative to a baseline, obtained from the same sets of simulations, when we always selected the first embryo. The baseline risk is indicated in each legend as “bl”. Note that the baseline risk depends on the variance explained by the PRS, because the parental PRSs are determined as percentiles of the population distribution of the score, which has variance 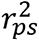. Finally, we computed the theoretical disease risk for the HRE strategy using Eq. (29) from *Materials and Methods*, the disease risk for the LRP strategy using Eq. (23), and the relative risk reduction (shown in curves) for both strategies using Eq. (36).

It is also conceivable that families may be more likely to seek PES when one or both prospective parents is affected by a given disease. In **Figure 3 - Figure Supplement 1**, we plot the RRR under the HRE and LRP strategies given that the parents are both healthy, both affected, or one of each (where we fixed the prevalence *K* = 5% and the heritability to ℎ^2^ = 40%). The figure illustrates that parental disease status has relatively little impact on the expected RRR (especially in comparison to the changes under HRE when conditioning on the actual parental PRSs). This is because, as long as 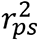 ≪ 1, parental disease does not necessarily provide much information about parental PRS, and thus does not strongly constrain the number of risk alleles available to each embryo.

### Absolute vs relative risk

The above results were presented in terms of *relative* risk reductions. However, **Figure 3 - Figure Supplement 1** also shows the baseline risk of an embryo of parents with a given disease status. For example, when one of the parents is affected, selecting the lowest risk embryo out of *n* = 5 (for a realistic 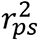= 0.1) reduces the risk from 10.0% to only 5.8%, thus nearly restoring the risk of the future child to the population prevalence (5%). More generally, we plot the *absolute* risk reduction (ARR) under the HRE and LRP strategies in **Figure 3 - Figure Supplement 2** for a few values of the parental PRSs. Notably, while RRRs under the LRP strategy somewhat decrease with increasing parental PRSs, the ARRs substantially increase, in accordance with an expectation that PES in higher-risk parents should eliminate more disease cases.

The clinical interpretation of these *absolute risk* changes will vary based on the population prevalence of the disorder (or the baseline risk of specific parents), and can offer a very different perspective on the magnitude of the effects (Gordis, 2014; Lázaro-Muñoz et al., 2020; Murray et al., 2020). In particular, for a *rare* disease, large *relative* risk reductions may result in very small changes in *absolute* risk. As an example, schizophrenia is a highly heritable (Sullivan et al., 2003) serious mental illness with prevalence of at most 1% (Perälä et al., 2007). The most recent large-scale GWAS meta-analysis for schizophrenia (Schizophrenia Working Group of the Psychiatric Genomics Consortium, 2020) has reported that a PRS accounts for approximately 8% of the variance on the liability scale. Our model shows that a 52% RRR is attainable using the LRP strategy with *n* = 5 embryos. However, this translates to only ≈0.5 percentage points reduction on the absolute scale: a randomly-selected embryo would have a 99% chance of not developing schizophrenia, compared to a 99.5% chance for an embryo selected according to LRP. In the case of a more common disease such as type 2 diabetes, with a lifetime prevalence in excess of 20% in the United States (Geiss et al., 2014), the RRR with *n* = 5 embryos (if the full SNP heritability of 17% (Y. Zhang et al., 2018) were achieved) is 43%, which would correspond to >8 percentage points reduction in absolute risk.

### Variability of the risk reduction across couples

The results depicted in Figure 2 describe the *average* risk reduction across the population, whereas the results in Figure 3 demonstrate results for specific combinations of parental risk scores. However, it remains unclear whether the large average risk reductions observed under the LRP strategy are driven by only a small proportion of couples. More generally, we would like to fully characterize the dependence of the risk reduction on parental PRSs, which could be of interest to physicians and couples in real-world settings.

To address these questions, we define a new risk reduction index, which we term the *per-couple* relative risk reduction, or pcRRR. Informally, the pcRRR is the relative risk reduction conditional of the PRSs of the couple. Mathematically, pcRRR(couple) 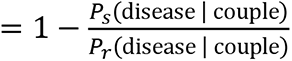. Here, *P_s_* (disease | couple) is the probability that the (PRS-based) *s*elected embryo is affected given the PRSs of the couple, and *P*_*r*_(disease | couple) is similarly defined for a *r*andomly selected embryo. Conveniently, the pcRRR depends only on the average of the maternal and paternal PRSs, which we denote as *c*. We calculated pcRRR(*c*) analytically under the LRP strategy (*Materials and Methods*), as well as computed the distribution of pcRRR(*c*) across all couples in the population.

We show the distribution of pcRRR(*c*) in **Figure 4**, panels (A)-(C). The results demonstrate that the pcRRR is relatively narrowly distributed around its mean, for all values of the prevalence (*K*) considered. The distribution becomes somewhat wider (and left-tailed) for the most extreme 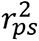 (0.3). Thus, the population-averaged RRRs are not driven by a small proportion of the couples. In agreement, the pcRRR depends only weakly on the average parental PRS, as can be seen in panels (D)-(F).

**Figure 4.**
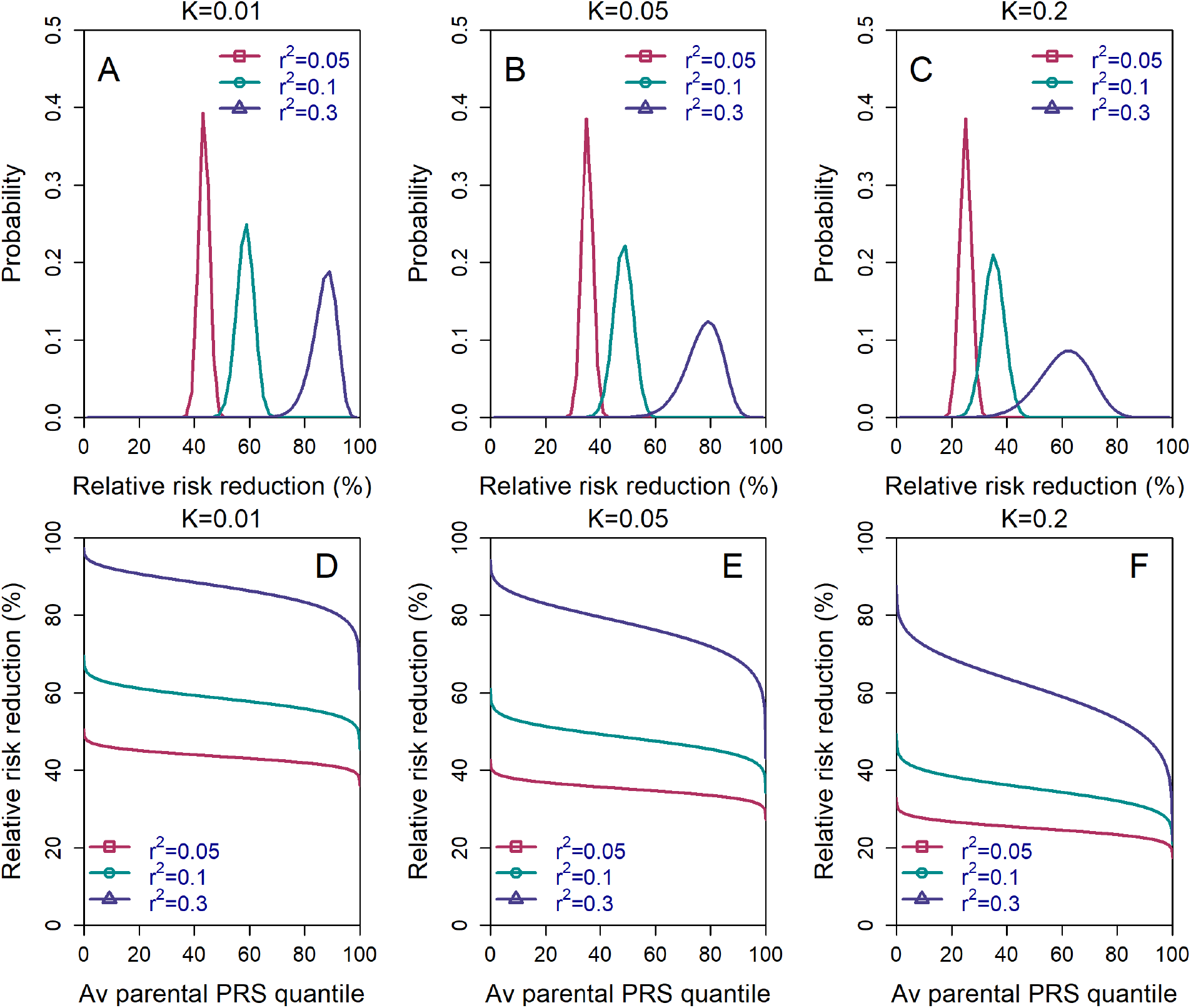
The variability in the relative risk reduction across couples. We considered only the *lowest-risk prioritization* strategy. In panels (A)-(C), we computed the theoretical distribution of the *per-couple* relative risk reduction, as explained in the *Materials and Methods*, *Appendix* Section 5. Briefly, the *per-couple* RRR is defined as 1 − *P*_*s*_ (disease | *c*)/*P*_*r*_(disease | *c*), where *P*_*s*_(disease | *c*) is the probability of an embryo *s*elected based on its PRS to be affected and *P*_*r*_ (disease | *c*) is the probability of a *r*andomly selected embryo to be affected. Our modeling suggests that *c*, which is the average of the paternal and maternal PRSs, is the only determinant of the relative risk reduction of a given couple. We computed the distribution of the *per-couple* RRR based on 10^4^ quantiles of *c*, thus covering all hypothetical couples in the population. The number of embryos was set to *n* = 5 in all panels. Panels (A)-(C) correspond to prevalence of *K* = 0.01, 0.05, and 0.02, respectively. In panels (D)-(F), we plot the theoretical RRR vs the quantile of the average parental PRS *c* (see *Materials and Methods* Section 5.1).

We note that the *per-couple* relative risk reduction is itself also an average, over all possible batches of *n* embryos of the couple. One may thus ask what is the distribution of possible RRRs across these batches. We provide a short discussion in *Materials and Methods* (*Appendix* Section 5.3).

### Pleiotropic effects of selection on genetically negatively correlated diseases

Polygenic risk scores are often correlated across diseases (Watanabe et al., 2019; Zheng et al., 2017). Therefore, selecting based on the PRS of one disease may increase or decrease risk for other diseases. While a full analysis of screening for multiple diseases is left for future work, our simulation framework allows us to investigate the potential harmful effects of prioritizing embryos for one disease, in case that disease is negatively correlated with another disease (*Materials and Methods*). We considered genetic correlations between diseases taking the values *ρ* = (−0.05, −0.1, −0.15, −0.2, −0.3). [The most negative correlation between two diseases reported in LDHub (http://ldsc.broadinstitute.org/ldhub/) is −0.3, occurring between ulcerative colitis and chronic kidney disease (Zheng et al., 2017).] In general, negative correlations between diseases are uncommon, and when they occur, typical correlations are about −0.1.

**Figure 5** shows the simulated risk reduction for the target disease and the risk increase for the correlated disease, across different values of *ρ* and for three values of the prevalence *K* (panels (A)-(C); assumed equal for the two diseases), all under the LRP strategy. In all panels, we used 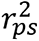 = 0.1 for both diseases. The relative risk reduction for the target disease is, as expected, always higher in absolute value than the risk increase of the correlated disease. For typical values of *ρ* = −0.1 and *n* = 5, the relative increase in risk of the correlated disease is relatively small, at ≈6% for *K* ≤ 0.05 and ≈3.5% for *K* = 0.2. However, for strong negative correlation (*ρ* = −0.3) the increase in risk can reach 22%, 16%, or 11% for *K* = 0.01, 0.05, and 0.2, respectively. Thus, care must be taken in the unique setting when the target disease is *strongly* negatively correlated with another disease.

**Figure 5.**
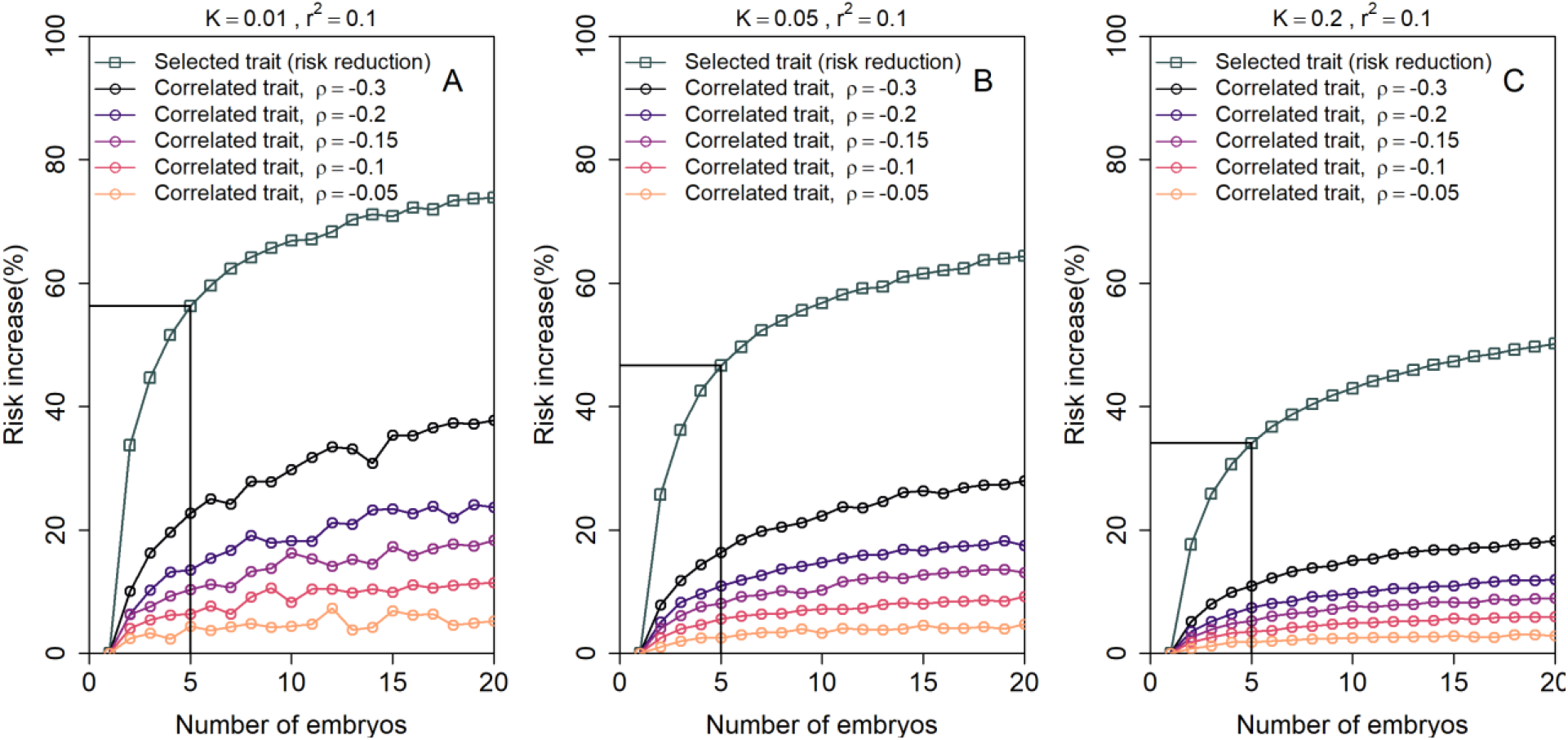
The increase in the risk of a negatively correlated disease due to polygenic embryo screening. We simulated two diseases that have genetic correlation *ρ* < 0. We assumed that the prevalence *K* is equal between the two diseases (*K* = 0.01, 0.05 and 0.2: panels (A)-(C), respectively), and that 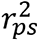 = 0.1 for both diseases. We simulated polygenic scores for the two diseases in *n* embryos in each of 10^6^ couples. For each couple, we selected the embryo either randomly or based on having the lowest PRS for the target disease. We then computed the risk of the embryo to have each disease as in the main analyses, by drawing the residual component of the liability and designating the embryo as affected if the total liability exceeded a threshold. The relative risk reduction of the target disease is shown as gray squares (and connecting lines) at the top of each plot. The relative risk *increase* for the correlated disease is shown in colored circles (and connecting lines), with different colors corresponding to different values of *ρ* (see legend). Note that the risk reduction for the target disease is independent of *ρ*.

### Simulations based on real genomes from case-control studies

Our analysis so far has been limited to mathematical analysis and simulations based on a statistical model. In principle, it would be desirable to compare our predictions to results based on real data. However, clearly, no real genomic and phenotypic data exist that would correspond to our setting, nor could such data be ethically or practically generated. Thus, we resort to a “hybrid” approach, in which we simulate the genomes of embryos based on real genomic data from case-control studies. This approach is similar to the one we have previously used for studying polygenic embryo screening for traits (Karavani et al., 2019).

Briefly, our approach is as follows. We consider separately two diseases with somewhat differing genetic architecture: schizophrenia, which is amongst the most polygenic complex diseases, with no common loci of high effect size, and Crohn’s disease, which is estimated to be less polygenic, and has several common loci with much larger effects than those found in schizophrenia (O’Connor et al., 2019). For each disease, we used genomes of unrelated individuals drawn from case-control studies. For schizophrenia, we used ≈900 cases and ≈1600 controls of Ashkenazi Jewish ancestry, while for Crohn’s, we used ≈150 cases and ≈100 controls of European ancestry. We then generated “virtual couples” by randomly mating pairs of individuals, regardless of sex or disease status. For each couple, we simulate the genomes of *n* hypothetical embryos, based on the laws of Mendelian inheritance and by randomly placing crossovers according to genetic map distances. In parallel, we used the “parental” genomes to learn a logistic regression model that predicts the disease risk given a PRS computed based on existing summary statistics. We then computed the PRS of each simulated embryo, and predicted the risk that embryo to be affected. Finally, we compared the risk of disease between a population in which one embryo per couple is selected at random, vs. a population in which one embryo is selected based on its PRS. For complete details, see *Materials and Methods*.

In **Figure 6**, we plot the results for the relative risk reduction for schizophrenia (panels (A) and (B)) and Crohn’s disease (panels (C) and (D)). For each disease, we consider both the HRE and LRP strategies. The analytical predictions closely match the empirical risk reductions generated in the simulations, except for a slight overestimation of the RRR under the LRP strategy. Nevertheless, for both schizophrenia and Crohn’s disease, we empirically observe that RRRs as high as ≈45% are achievable with *n* = 5 embryos. In contrast, under the HRE strategy and when excluding embryos at the top 2% risk percentiles, risk reductions are very small, in agreement with the theoretical predictions. These results thus provide support to the robustness of our statistical model.

**Figure 6.**
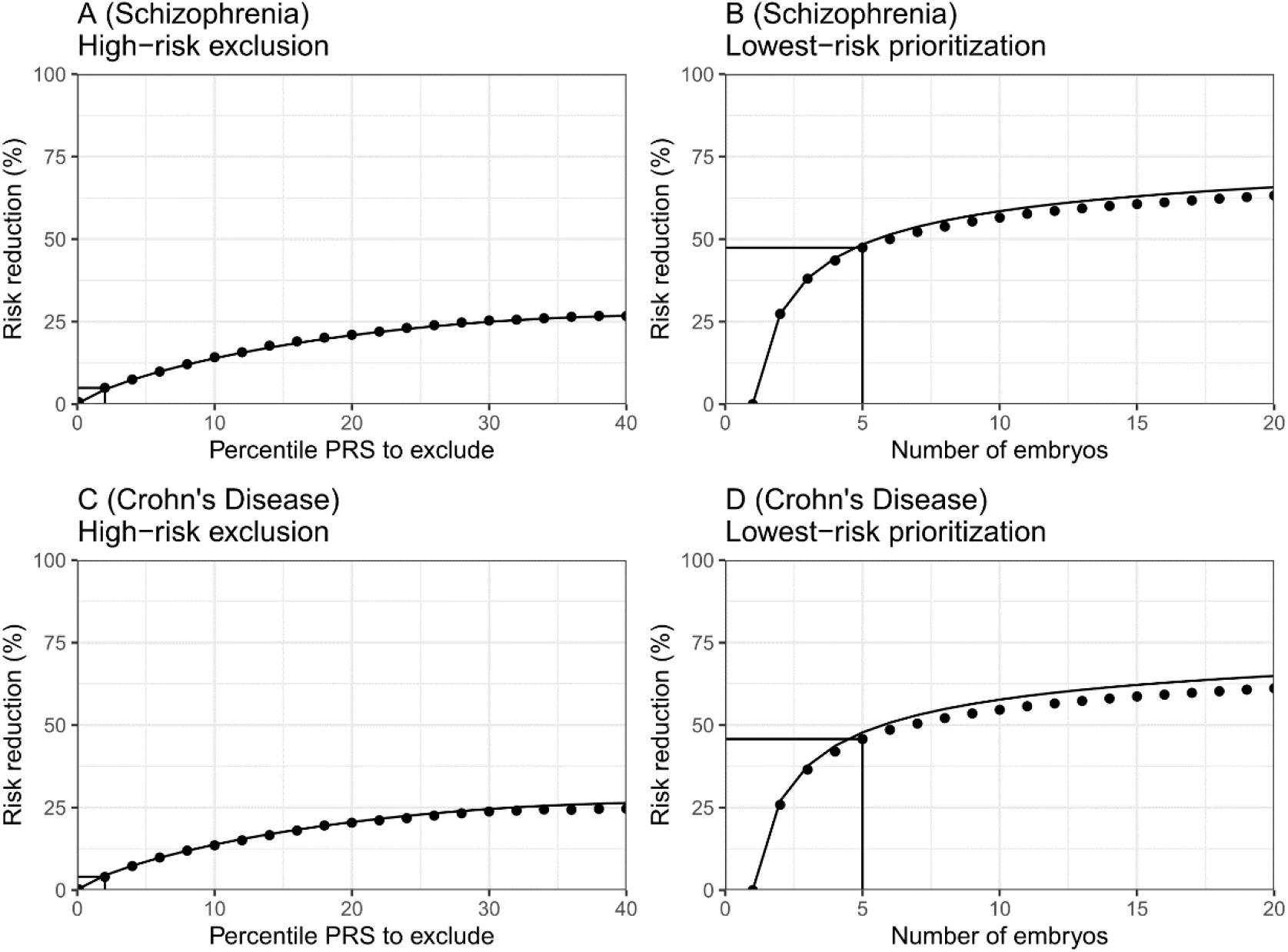
The empirical relative risk reduction in simulated embryos based on genomes from case-control studies of schizophrenia and Crohn’s disease. We used ≈900 cases and ≈1600 controls for schizophrenia, and ≈150 cases and ≈100 controls for Crohn’s. For each disease, we drew 5,000 random “virtual couples”, regardless of sex, but correcting for case/control ascertainment. For each such random couple, we simulated the genomes of up to *n* = 20 embryos (children) based on Mendelian segregation and published recombination maps. For each embryo, we computed the PRS for the given disease (schizophrenia or Crohn’s) using the most recent summary statistics that exclude our cohort. We computed the risk of each embryo to be affected based on a logistic regression model we learned in the “parental” cohort. Panels (A) and (B) show results for schizophrenia, while panels (C) and (D) show results for Crohn’s. In panels (A) and (C), we plot the relative risk reduction (RRR) under the *high-risk exclusion* (HRE) selection strategy, in which an embryo was randomly selected (out of *n* = 5 embryos), unless its PRS was above a given percentile. The RRR was computed against a baseline strategy of selection of an embryo at random, and is plotted vs the exclusion percentile. In panels (B) and (D), we show the relative risk reduction under the *lowest-risk prioritization* (LRP) strategy, in which the embryo with the lowest PRS was selected. We plot the RRR vs the number of embryos *n*. In all panels, dots correspond to the results of simulations, and solid lines correspond to the theory. The theory was computed assuming prevalence of 1% for schizophrenia and 0.5% for Crohn’s, and variance explained on the liability scale of 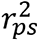 = 0.068 for schizophrenia 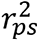 = 0.056 for Crohn’s (calculated using the method of (Lee et al., 2012)). Further details are provided in *Materials and Methods*.

To further investigate the assumptions of our model, we test in **Figure 6 – Figure Supplement 1** two intermediate predictions. The first is that the variance of the PRSs of embryos of a given couple should not depend on the average parental PRS. This is indeed the case (panels (A) and (C)), with the only exception of an uptick of the variance at very low parental PRSs for schizophrenia. The second prediction is that the variance across embryos is half of the variance in the parental population. The empirical results again show reasonable agreement with the theoretical prediction (panels (B) and (D)). The empirical variance (averaged across couples) was slightly lower than expected (by ≈4% for schizophrenia and ≈14% for Crohn’s), which may explain our slight overestimation of the expected RRR under the LRP strategy.

## Discussion

In this paper, we used statistical modeling to evaluate the expected outcomes of screening embryos based on polygenic risk scores for a single disease. We predicted the relative and absolute risk reductions, either at the population level or at the level of individual couples. Our model is flexible, allowing us to provide predictions across various values of, e.g., the PRS strength, the disease prevalence, the parental PRS or disease status, and the number of available embryos. We presented a comprehensive analysis of the expected outcomes across various settings, including when there is a concern about a second disease negatively correlated with the target disease. We finally validated our modeling assumptions using genomes from case-control studies. Our publicly available code could help researchers and other stakeholders estimate the expected outcomes for settings we did not cover.

Our most notable result was that a crucial determinant of risk reduction is the *selection strategy*. The use of PRS in adults has focused on those at highest risk (Chatterjee et al., 2016; Dai et al., 2019; Gibson, 2019; Khera et al., 2018; Mars et al., 2020; Mavaddat et al., 2019; Torkamani et al., 2018), for whom there may be maximal clinical benefit of screening and intervention. However, as PRSs have relatively low sensitivity, such a strategy is relatively ineffective in reducing the overall population disease burden (Ala-Korpela & Holmes, 2020; Wald & Old, 2019). Similarly, in the context of PES, exclusion of high-risk embryos will result in relatively modest risk reductions. By contrast, selecting the embryo with the lowest PRS may result in large reductions in relative risk.

While our prior work (Karavani et al. 2019) demonstrated that PES would have a small effect on quantitative traits, here we show that a small reduction in the liability can lead to a large reduction in the proportion of affected individuals. This is fundamentally a property of a threshold character with an underlying normally distributed continuous liability. For such traits, most of the individuals in the extreme of the liability distribution (i.e., the ones affected) are concentrated very near the threshold. Thus, even slightly reducing their liability can move a large proportion of affected individuals below the disease threshold. However, it should be noted that conventional thresholds for defining presence of disease may contain some degree of arbitrariness if the underlying distribution of pathophysiology is truly continuous. Consequently, the effects on ultimate morbidity may depend on the validity of the threshold itself (Davidson & Kahn, 2016).

We investigated how the range of potential PES outcomes varies with the PRSs of the parents or with their disease status. Under the HRE strategy, if only excluding embryos at the few topmost risk percentiles, the RRR is very small when the parents have low PRSs, and vice versa (Figure 3, panels (A)- (D)). This is expected, as excluding high PRS embryos will be effective only for couples who are likely to have many such embryos. Under the LRP strategy, the RRR depends only weakly on the parental PRSs (Figure 3, panels (E)-(H), and Figure 4). Under both strategies, the relative risk reduction depends only weakly on the parental disease status, as parental disease status is a weak signal for the underlying PRS. However, the *absolute* risk reduction increases substantially with increasing parental PRSs (**Figure 3 – Figure Supplement 2**) and when one or more parents are affected.

Our study has several limitations. First, our results assume an infinitesimal genetic architecture for the disease, which may not be appropriate for oligogenic diseases and is not relevant for monogenic disorders. However, it has been repeatedly demonstrated that common, complex traits and diseases are highly polygenic (Gazal et al., 2017; Holland et al., 2020; O’Connor et al., 2019; Shi et al., 2016; Zeng et al., 2018, 2021). For example, it was recently estimated that for almost all traits and diseases examined, the number of independently associated loci was at least ≈350, reaching ≈10,000 or more for cognitive and psychiatric phenotypes (O’Connor et al., 2019). This provides more than sufficient variability for the PRS to attain a normal distribution in the population and for our modeling assumptions to hold. Indeed, our empirical results for schizophrenia and Crohn’s disease, two diseases with somewhat different genetic architectures, agreed reasonably well with the theoretical predictions. However, our models would need to be substantially adjusted in the presence of variants of very large effect, such as inherited or *de novo* coding variants or copy number variants, e.g., as in autism (Satterstrom et al., 2020; Takumi & Tamada, 2018).

Additionally, our model relies on several simplifying statistical assumptions. For example, we did not explicitly model assortative mating, although this seems reasonable given that for genetic disease risk, correlation between parents is weak (Rawlik et al., 2019), and given that our previous study of traits showed no difference in the results between real and random couples (Karavani et al., 2019). This deficiency is also partly ameliorated by our modeling of the risk reduction when explicitly given the parental PRSs or disease status. Another assumption we made is that environmental influences on the child’s phenotype are independent of those that have influenced the parents (when conditioning on the parental disease status). However, this is reasonable given that family-specific environmental effect have been shown to be weak for complex diseases (Wang et al., 2017). For a discussion of additional model assumptions, see *Materials and Methods*, *Appendix* section 10.

Perhaps more importantly, we assumed throughout that 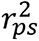 represents the realistic accuracy of the PRS achievable, within-family, in a real-world setting in the target population. However, the realistically achievable 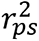 may be lower than reported in the original publications that have generated the scores. For example, the accuracy of PRSs is sub-optimal when applied in non-European populations and across different socio-economic groups (Duncan et al., 2019; Mostafavi et al., 2020). A PRS that was tested on adults may be less accurate in the next generation. Additionally, the variance explained by the score, as estimated in samples of unrelated individuals, is inflated due to population stratification, assortative mating, and indirect parental effects (Kong et al., 2018; Young et al., 2019; Morris et al., 2020; Mostafavi et al., 2020). The latter, also called “genetic nurture”, refers to trait-modifying environmental effects induced by the parents based on their genotypes. These effects do not contribute to prediction accuracy when comparing polygenic scores between siblings (as when screening IVF embryos), and thus, the variance explained by polygenic scores in this setting can be substantially reduced, in particular for cognitive and behavioral traits (Howe et al., 2021; Selzam et al., 2019). Our risk reduction estimates thus represent an upper bound relative to real-world scenarios. On the other hand, recent empirical work on within-family disease risk prediction showed that the reduction in accuracy is at most modest (Lello et al., 2020), and within-siblings-GWAS yielded similar results to unrelated-GWAS for most physiological traits (Howe et al., 2021). Additionally, accuracy in non-European populations is rapidly improving due to the establishment of national biobanks in non-European countries (Koyama et al., 2020; Vujkovic et al., 2020) and improvement in methods for transferring scores into non-European populations (Amariuta et al., 2020; Cai et al., 2021). Either way, the analytical results presented in this paper are formulated generally as a function of the achievable accuracy 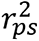, and as such, users can substitute values relevant to their specific target population and disease.

Another major limitation of this work is that we have only considered screening for a single disease. In reality, couples may seek to profile an embryo on the basis of multiple disease PRSs simultaneously, or based a global measure of lifespan or healthspan (Sakaue et al., 2020; Timmers et al., 2020; Zenin et al., 2019). This is likely to reduce the per-disease risk reduction, as we have previously observed for quantitative traits (Karavani et al., 2019), but will also likely be more cost effective (Treff et al., 2020). PES for multiple diseases requires the formulation and analysis of new selection strategies, and is substantially more mathematically complex; we therefore leave it for future studies.

As our approach was statistical in nature, it is important to place our results in the context of real-world clinical practice of assisted reproductive technology. The number of embryos utilized in the calculations in the present study refers to viable embryos that could lead to *live birth*, which can be substantially smaller than the raw number of fertilized oocytes or even the number of implantable embryos at day 5. This consideration is especially important given the steep drop in risk reduction when the number of available embryos drops below 5 (**Figure 2**). In fact, many IVF cycles do not achieve *any* live birth. Rates of live birth decline with maternal age, in particular after age 40 (Smith et al., 2015); for women age >42, fewer than 4% of IVF cycles result in live births, making PES impractical. On the other hand, success rates will likely be higher for young prospective parents who seek PES to reduce disease risk but do not suffer from infertility. However, the prospect of elective IVF for the purpose of PES in such couples must be weighed against the potential risks of these invasive procedures to the mother and child (Dayan et al., 2019; Luke, 2017).

A different concern is whether the embryo biopsy (which is required for genotyping) may cause risk to the viability and future health of the embryo. Several recent studies have demonstrated no evidence for potential adverse effects of trophectoderm biopsy on rates of successful implantation, fetal anomalies, and live birth (Awadalla et al., 2021; He et al., 2019; Riestenberg et al., 2021; Tiegs et al., 2021). Moreover, no significant adverse effects have been detected for postnatal child development in a recent meta-analysis (Natsuaki & Dimler, 2018). On the other hand, a number of studies have reported that trophectoderm biopsy was associated with pregnancy complications, including preterm birth, pre-eclampsia, and hypertensive disorders of pregnancy (Li et al., 2021)(W. Y. Zhang et al., 2019)(Makhijani et al., 2021). Specific variations in biopsy protocols may account for differences in outcomes across studies (Rubino et al., 2020). Newly developed techniques may allow in the future to genotype an embryo non-invasively based on DNA present in spent culture medium, although the accuracy of these methods is still being debated (Leaver & Wells, 2020). It should also be noted that, throughout this manuscript, we assumed the use of single embryo transfer.

Finally, the results of our study invite a debate regarding ethical and social implications. For example, the differential performance of PES across selection strategies and risk reduction metrics may be difficult to communicate to couples seeking assisted reproductive technologies (Cunningham et al., 2015; Wilkinson et al., 2019). Indeed, in the first PES case report, the couple elected to forego any implantation despite the availability of embryos that were designated as normal risk (Treff et al. 2019). These difficulties are expected to exacerbate the already profound ethical issues raised by PES (as we have recently reviewed (Lázaro-Muñoz et al., 2020)), which include stigmatization (McCabe & McCabe, 2011), autonomy (including “choice overload” (Hadar & Sood, 2014)), and equity (Sueoka, 2016). In addition, the ever-present specter of eugenics (Lombardo, 2018) may be especially salient in the context of the LRP strategy. How to juxtapose these difficulties with the potential public health benefits of PES is an open question. We thus call for urgent deliberations amongst key stakeholders (including researchers, clinicians, and patients) to address governance of PES and for the development of policy statements by professional societies. We hope that our statistical framework can provide an empirical foundation for these critical ethical and policy deliberations.

## Materials and Methods

### Summary of the modeling results

In this section, we provide a brief overview of our model and derivations, with complete details appearing in the *Appendix*.

Our model is follows. We write the polygenic risk scores of a batch of *n* IVF embryos as (*s*_1_, …, *s*_*n*_), and generate the scores as *s*_*i*_ = *x*_*i*_ + *c*. The (*x*_1_, …, *x*_*n*_) are embryo-specific independent random variables with distribution 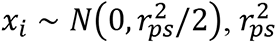 is the proportion of variance in liability explained by the score, and *c* is a shared component with distribution 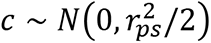, also representing the average of the maternal and paternal scores.

In each batch, an embryo is selected according to the selection strategy. Under *high-risk exclusion*, we select a random embryo with score *s* < *z*_*q*_*r*_*ps*_, where *z*_*q*_ is the (1 − *q*)-quantile of the standard normal distribution. If no such embryo exists, we select a random embryo, but we also studied the rule when in such a case, the lowest scoring embryo is selected. Under *lowest-risk prioritization*, we select the embryo with the lowest value of *s*. We computed the liability of the selected embryo as *y* = *s* + *e*, where 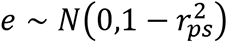. We designate the embryo as affected if *y* > *z_K_*, where *z_K_* is the (1 − *K*)-quantile of the standard normal distribution and *K* is the disease prevalence. In the simulations, we computed the disease probability (for each parameter setting) as the fraction of batches (out of 10^6^ repeats) in which the selected embryo was affected. We also simulated the score and disease status of a second disease, which is not used for selecting the embryo, but may be negatively correlated with the target disease.

We computed the disease probability analytically using the following approaches. We first computed the distribution of the score of the selected embryo. For *lowest-risk prioritization*, we used the theory of order statistics. For *high-risk exclusion*, we first conditioned on the shared component *c*, and then studied separately the case when all embryos are high-risk (i.e., have score *s* > *z*_*q*_*r*_*ps*_), in which the distribution of the unique component of the selected embryo (*x*) is a normal variable truncated from below at *z*_*q*_*r*_*ps*_, and the case when at least one embryo has score *s* < *z*_*q*_*r*_*ps*_, in which *x* is a normal variable truncated from above. We then integrated over the non-score liability components (and over *c* in some of the settings) in order to obtain the probability of being affected. We solved the integrals in the final expressions numerically in R.

We computed the risk reduction based on the ratio between the risk of a child of a random couple when the embryo was selected by PRS and the population prevalence. We also provide explicit results for the case when the average parental PRS *c* is known. These expressions allowed us to compute the distribution of risk reductions *per-couple*. Finally, when conditioning on the parental disease status, we integrated the disease probability of the selected embryo over the posterior distribution of the parental score and non-score genetic components. For full details and for an additional discussion of previous work and limitations, see the *Appendix*. R code is available at: https://github.com/scarmi/embryo_selection.

### Simulations based on genomes from case-control studies

Our main analysis has been limited to mathematical modeling of polygenic scores and their relation to disease risk. For obvious ethical and practical reasons, we could not validate our modeling predictions with actual experiments. Nevertheless, we could perform realistic simulations based on genomes from case-control studies, similarly to our previous work (Karavani et al., 2019). Our approach is generally as follows. We consider, separately, two diseases: schizophrenia and Crohn’s. For schizophrenia, we use ≈900 cases and ≈1600 controls of Ashkenazi Jewish ancestry, while for Crohn’s, we use ≈150 cases and ≈100 controls from the New York area. For each disease, we use these individuals, who are unrelated, to generate “virtual couples” by randomly mating pairs of individuals. For each such “couple”, we simulate the genomes of *n* hypothetical embryos, based on the laws of Mendelian inheritance and by randomly placing crossovers according to genetic map distances. In parallel, we use the same genomes to learn a logistic regression model that predicts the risk of disease given a PRS computed from the most recently available summary statistics (excluding the samples in our test cohorts). We then compute the PRS of each simulated embryo, and predict the risk of disease of that embryo. We finally compare the risk of disease between one randomly selected embryo per couple vs one embryo selected based on PRS. In the paragraphs below, we provide additional details.

#### The Ashkenazi schizophrenia cohort

The samples and the genotyping process were previously described (Lencz et al., 2013). Patients were recruited from hospitalized inpatients at seven medical centres in Israel diagnosed with schizophrenia or schizoaffective disorder and samples from healthy Ashkenazi individuals were collected from volunteers at the Israeli Blood Bank. All subjects provided written informed consent, and corresponding institutional review boards and the National Genetic Committee of the Israeli Ministry of Health approved the studies. DNA was extracted from whole blood and genotyped for ∼1 million genome-wide SNPs using Illumina HumanOmni1-Quad arrays. We performed the following quality control steps. First, we removed samples with (1) genotyping call rate <95%; (2) one of each pair of related individuals (total shared identical-by-descent (IBD) segments >700cM); and (3) sharing of less than 15cM on average with the rest of the cohort (indicating non-Ashkenazi ancestry). We removed SNPs with (1) call rate <97%; (2) minor allele frequency <1%; (3) significantly different allele frequencies between males and females (P-value threshold = 0.05/#SNPs); (4) differential missingness between males and females (P<10^-7^) based on a χ^2^ test; (5) deviations from Hardy-Weinberg equilibrium in females (P-value threshold = 0.05/#SNPs); (6) SNPs in the HLA region (chr6:24-37M); and (7) (after phasing) SNPs having A/T or C/G polymorphism, as we could not unambiguously link them to corresponding effect sizes in the summary statistics. We finally used autosomal SNPs only. The remaining number of individuals was 2,526 (897 cases and 1629 controls), and the number of SNPs was 728,505. We phased the genomes using SHAPEIT v2 (Delaneau et al., 2013).

#### The Mt Sinai Crohn’s disease cohort

Samples from subjects with Crohn’s disease were recruited from clinics by Mt Sinai providers. All subjects provided written, informed consent in studies approved by the Mt Sinai Institutional Review Board. Genotyping was performed at the Broad Institute using the Illumina Global Screening Array (GSA) chip, as previously described (Gettler et al., 2021). We phased the genomes using Eagle v2.4.1 (Loh et al., 2016). We then removed SNPs having A/T or C/G polymorphism. The remaining number of individuals was 257 (154 cases and 103 controls) and the number of SNPs was 560,612.

#### Simulating couples and embryos

For each disease, we generated 5,000 unique couples by randomly pairing individuals (regardless of their sex) according to the population prevalence of the disease. For example, for schizophrenia, assuming a prevalence of 1%, a proportion 0.99^2^ of the couples were both controls. Given a pair of parents, we simulated 20 offspring (embryos) by specifying the locations of crossovers in each parent. Recombination was modeled as a Poisson process along the genome, with distances measured in cM using sex-averaged genetic maps (Bhérer et al., 2017). Specifically, for each parent and embryo, we drew the number of crossovers in each chromosome from a Poisson distribution with mean equal to the chromosome length in Morgan. We then determined the locations of the crossovers by randomly drawing positions along the chromosome (in Morgan). We mixed the phased paternal and maternal chromosomes of the parent according to the crossover locations, and randomly chose one of the resulting sequences as the chromosome to be transmitted to the embryo. We repeated for the other parent, in order to form the diploid genome of the embryo.

#### Developing a polygenic risk score for schizophrenia

We used summary statistics from the most recent schizophrenia GWAS of the Psychiatric Genomics Consortium (PGC) (Schizophrenia Working Group of the Psychiatric Genomics et al., 2020). Note that we specifically used summary statistics that excluded our Ashkenazi cohort. We used the entire cohort (2526 individuals) to estimate linkage disequilibrium (LD) between SNPs, and performed LD-clumping on the summary statistics in PLINK (Chang et al., 2015), with a window size of 250kb, a minimum *r*^2^ threshold for clumping of 0.1, a minimum minor allele frequency threshold of 0.01, and a maximum P-value threshold of 0.05. The P-value threshold was chosen based on results from the PGC study. After clumping, the final score included 23,036 SNPs. To construct the score, we used the effect sizes reported in the GWAS summary statistics, without additional processing.

#### Developing a polygenic risk score for Crohn’s disease

We used summary statistics derived from European samples available from https://www.ibdgenetics.org/downloads.html (Liu et al., 2015), which did not include our cohort. We estimated LD using the entire Crohn’s disease cohort, and performed LD-clumping and P-value thresholding using the same parameters as for the schizophrenia cohort, as described above. The final score included 9,403 SNPs.

#### Calculating the PRS and the risk of an embryo

For each disease, we calculated polygenic scores for each parent and simulated embryo in PLINK, using the --score command with default parameters. Using the polygenic scores of the parents, we fitted a logistic regression model for the case/control status as a function of the polygenic scores. We did not adjust for additional covariates: for schizophrenia, genetic ancestry is homogeneous in our Ashkenazi cohort, and age and sex contributed very little to predictive power (increased AUC from 0.695 only to 0.717). For Crohn’s, age was not available, and sex did not contribute to predictive power (increased AUC from to 0.693 to 0.695). We adjusted the intercept of the logistic regression models to account for the case-control sampling (Rose & van der Laan, 2008). We then used the model to predict the probability that a simulated embryo would develop the disease.

To determine the percentiles of the PRS for each disease, we derived an approximation to the distribution of the PRS in the population by fitting a normal distribution to the scores in our dataset. To take into account the case/control ascertainment, we weighted the case and control samples according to the population prevalence of the disease (1% for schizophrenia (Perälä et al., 2007) and 0.5% for Crohn’s (GBD 2017 Inflammatory Bowel Disease Collaborators, 2020). We calculated the weighted mean and variance of the scores using the wtd.mean and wtd.var functions in the HMisc package in R. A normal distribution with the resulting mean and variance was used to calculate percentiles of the scores. The percentiles were then used to select (simulated) embryos under the *high-risk exclusion* strategy (see below).

#### Calculating the risk reduction

For each disease, we performed the following simulations. For each selection strategy (either *high-risk exclusion* or *lowest-risk prioritization*), we selected one embryo for each couple according to the strategy, and computed the probability of disease for the selected embryo. We then averaged the risk over all couples. We similarly computed the risk under selection of a random embryo for each couple. We computed the relative risk reduction based on the ratio between the risk under PRS-based selection and the risk under random selection. To compare to the theoretical expectations, we estimated the variance explained by the score on the liability scale using the method of Lee et al. (Lee et al., 2012). Specifically, we first computed the correlation between the observed case/control status (coded as 1 and 0, respectively) and the PRS, and then used Eq. (15) in Lee et al to convert the squared correlation to the variance explained. We obtained 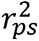 = 6.8% for schizophrenia, which is close to the 7.7% reported in the original GWAS paper (Schizophrenia Working Group of the Psychiatric Genomics et al., 2020), and 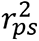 = 5.6% for Crohn’s disease. We then substituted this value and prevalence of *K* = 0.01 for schizophrenia and *K* = 0.005 for Crohn’s in our formulas for the relative risk reduction.

#### Code availability

Code for these analyses is at https://github.com/dbackenroth/embryo_selection.

## Abbreviations

IVF: *in-vitro* fertilization
PRS: polygenic risk score
PES: polygenic embryo screening
RRR: relative risk reduction
ARR: absolute risk reduction
HRE: high-risk exclusion
LRP: lowest-risk prioritization
LTM: liability threshold model
pcRRR: per-couple relative risk reduction.

## Acknowledgements

We thank Gabriel Lázaro-Muñoz, Stacey Pereira, Chaim Jalas, and David A. Zeevi for helpful discussions.

---

**Figure 2 - Figure Supplement 1.**
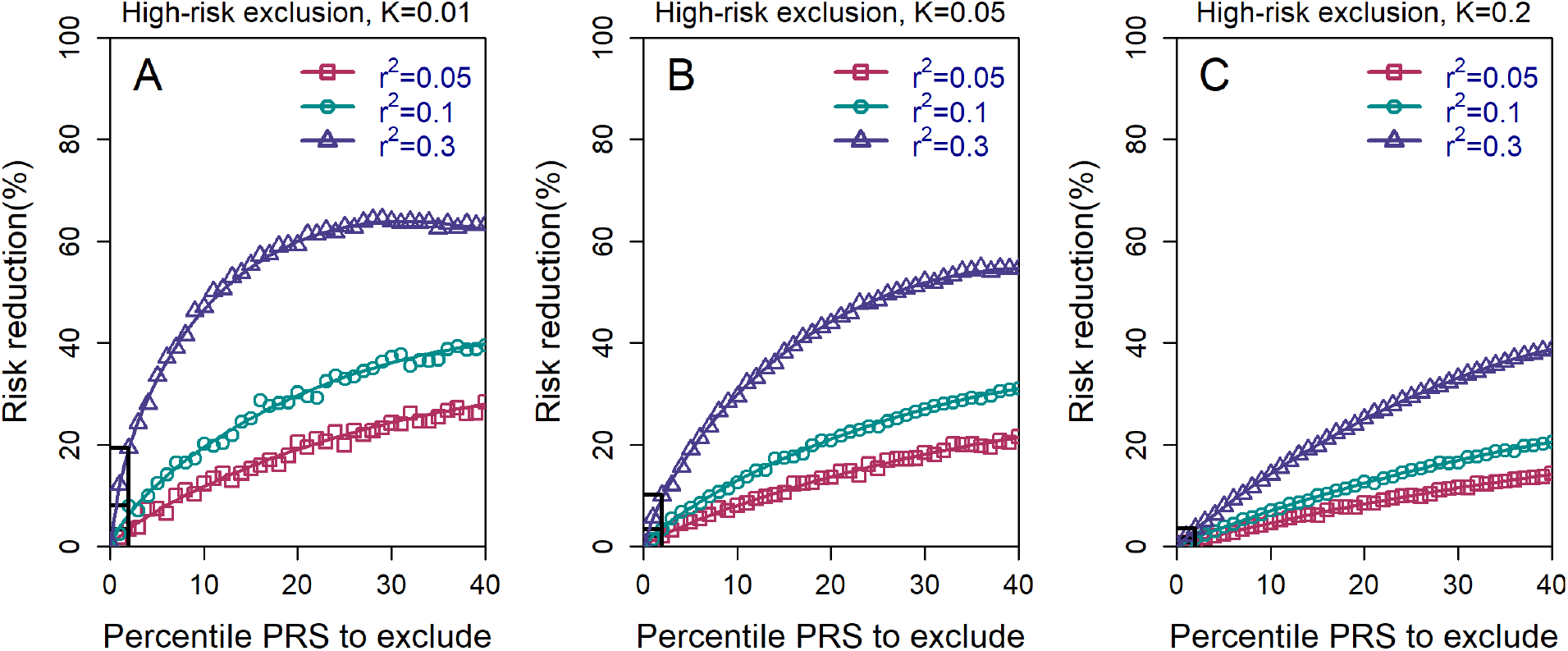
The relative risk reduction for the *high-risk exclusion* strategy, with *n* = 10 available embryos. All details are exactly as in panels (A-C) in **Figure 2** of the main text, except that we simulated *n* = 10 embryos.

**Figure 2 – Figure Supplement 2.**
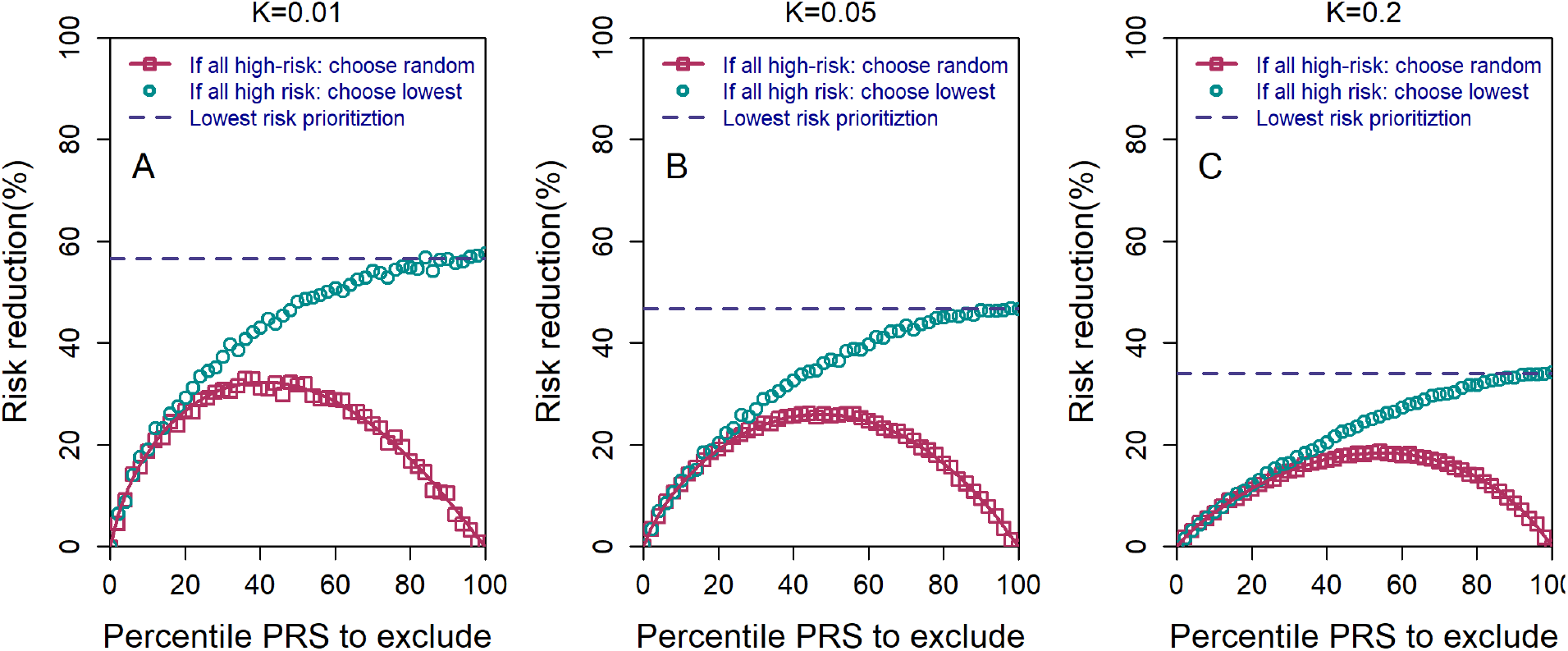
The relative risk reduction under the high-risk exclusion (HRE) strategy, using two different rules for how an embryo is selected when all embryos are high risk. All details are similar to those of Figure 2 of the main text, except the following. We used *n* = 5 embryos, 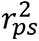 = 0.1, and *K* = 0.01, 0.05, and 0.2 (panels (A)-(C), respectively). For both sub-strategies, we first determined whether there were any non-high-risk embryos. If such embryos existed, one of them was randomly selected. If all embryos were high risk, the pink symbols and lines correspond to selecting an embryo at random (symbols: simulations; line: theory; see *Materials and Methods*). The cyan symbols correspond to selecting the embryo with the lowest PRS (simulations only). The blue dashed horizontal line corresponds to the theoretical relative risk reduction for the *lowest-risk prioritization* (LRP) strategy. When all embryos are designated as high risk (percentile PRS to exclude is 100%), the random selection sub-strategy reduces to a completely random selection and thus yields no risk reduction, whereas the lowest PRS sub-strategy becomes equivalent to the regular LRP strategy.

**Figure 2 – Figure Supplement 3.**
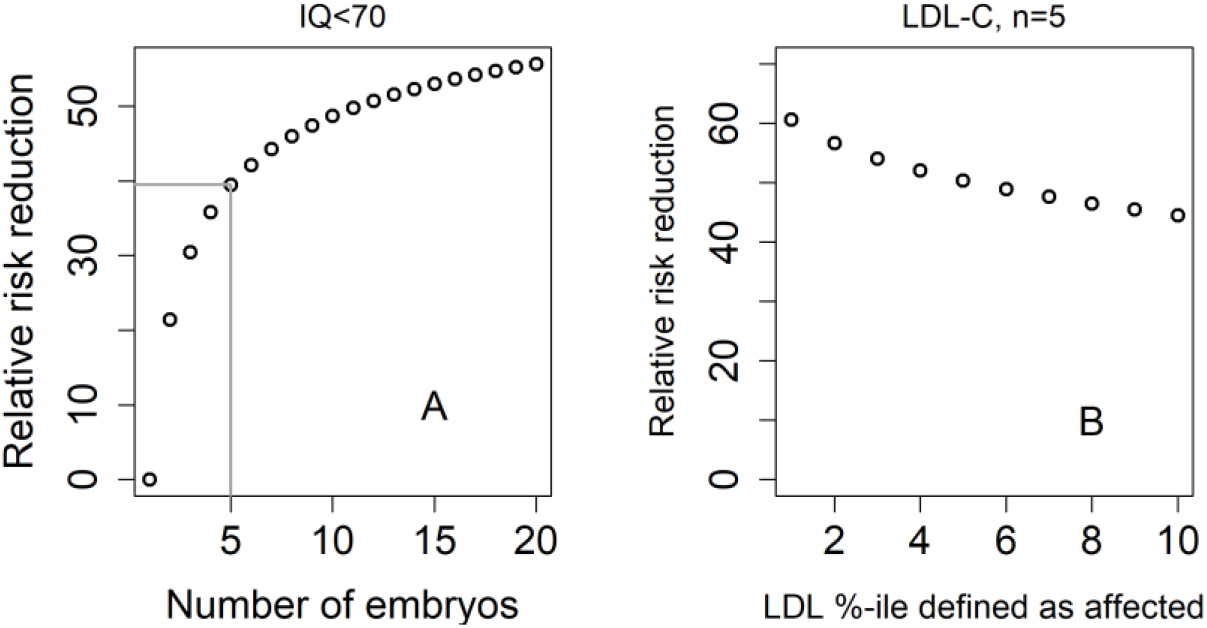
The relative risk reduction under the *lowest-risk prioritization* strategy for a dichotomized trait. In panel (A), we define a hypothetical individual as “affected” (or having an intellectual disability) if that individual has IQ<70. Assuming IQ is normally distributed with a mean of 100 and a SD of 15, this implies that the prevalence is *K* = 2.3%. We plot the predicted relative risk reduction (computed as in Figure 2) vs the number of embryos *n* under the LRP strategy (note that here, the embryo with the *highest* score is selected). We used 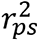 = 0.052 (Savage et al., 2018). In panel (B), we show the predicted RRR under the LRP strategy for a “high LDL cholesterol” binary trait. Here, we fixed *n* = 5 and varied the prevalence of our hypothetical “high LDL” trait. Given a prevalence *K*, an individual is defined as having “high LDL” if its LDL value in the top *K*-percentiles. We used 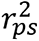 = 0.12 (Weissbrod et al., 2021).

**Figure 3 - Figure Supplement 1.**
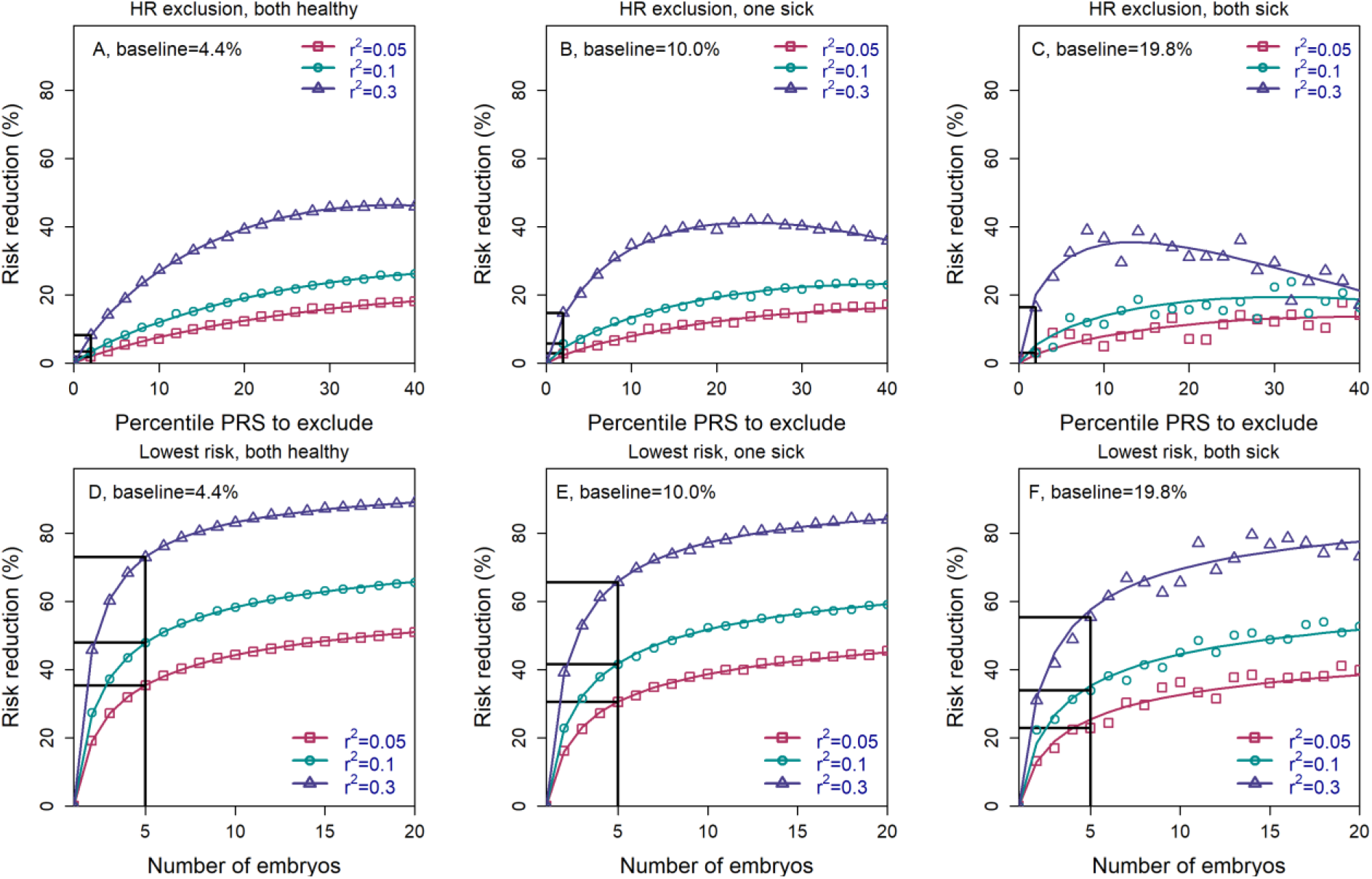
The relative risk reduction when the parental disease status is known. Panels (A)-(C) are for the *high-risk exclusion* (HRE) strategy, while panels (D)-(F) are for the *lowest-risk prioritization* (LRP) strategy. The details are as in Figure 2, except the following. First, we fixed the prevalence to *K* = 5% and the heritability to ℎ^2^ = 0.4 (note that the heritability was not needed in previous figures). Second, in the simulations, we first drew the parental genetic components: *s*_*m*_ and *w*_*m*_ for the mother, and *s*_*f*_ and *w*_*f*_ for the father, where *s*_*m*_ ∼ *s*_*f*_ ∼ *N*(0, 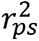) are the polygenic scores and *w*_*m*_ ∼ *w*_*f*_ ∼ *N*(0, ℎ^2^ − 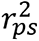) represent non-score genetic factors (*Materials and Methods*). We drew the environmental component for each parent as *ϵ*_*m*_ ∼ *ϵ*_*f*_ ∼ *N*(0,1 − ℎ^2^) and computed the liability of each parent as *s* + *w* + *ϵ*. If the liability of a parent exceeded *z*_*K*_ (the upper *K*-quantile of the standard normal distribution), we designated that parent as affected. We then stratified the risk reduction results based on the number of affected parents: 0 (panels (A) and (D), 1 (panels (B) and (E)), and 2 (panels (C) and (F)). Note that as expected, the number of families in which both parents are affected is small, and thus, the results in panels (C) and (F) are noisy. For each set of parents, we drew the PRS of each embryo as *s*_*i*_ = (*s*_*m*_ + *s*_*f*_)/2 + *x*_*i*_ (*i* = 1, …, *n*), where *x*_*i*_ ∼ *N*(0, 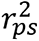 /2) is an embryo-specific component of the score (independent across embryos). We then selected one embryo from each family based on either selection strategy. We computed the liability of the selected embryo as *s*_*i*_ + (*w*_*m*_ + *w*_*f*_)/2 + *v*_*i*_ + *ϵ*_*i*_, where *v*_*i*_ ∼ *N*(0, (ℎ^2^ − 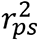)/2) is the embryo-specific component of the non-score genetic factors, and *ϵ*_*i*_ ∼ *N*(0,1 − ℎ^2^) is the environmental component of the embryo (*Materials and Methods*). The embryo was designated as affected or unaffected as described above for the parents. We computed the risk reduction (according to either strategy) relative to a baseline, obtained from the same sets of simulations when we always selected the first embryo. The baseline risk is indicated on top of each panel. We computed the theoretical relative risk reduction for the two strategies as summarized in *Appendix* Section 6.9 of the *Materials and Methods*.

**Figure 3 - Figure Supplement 2.**
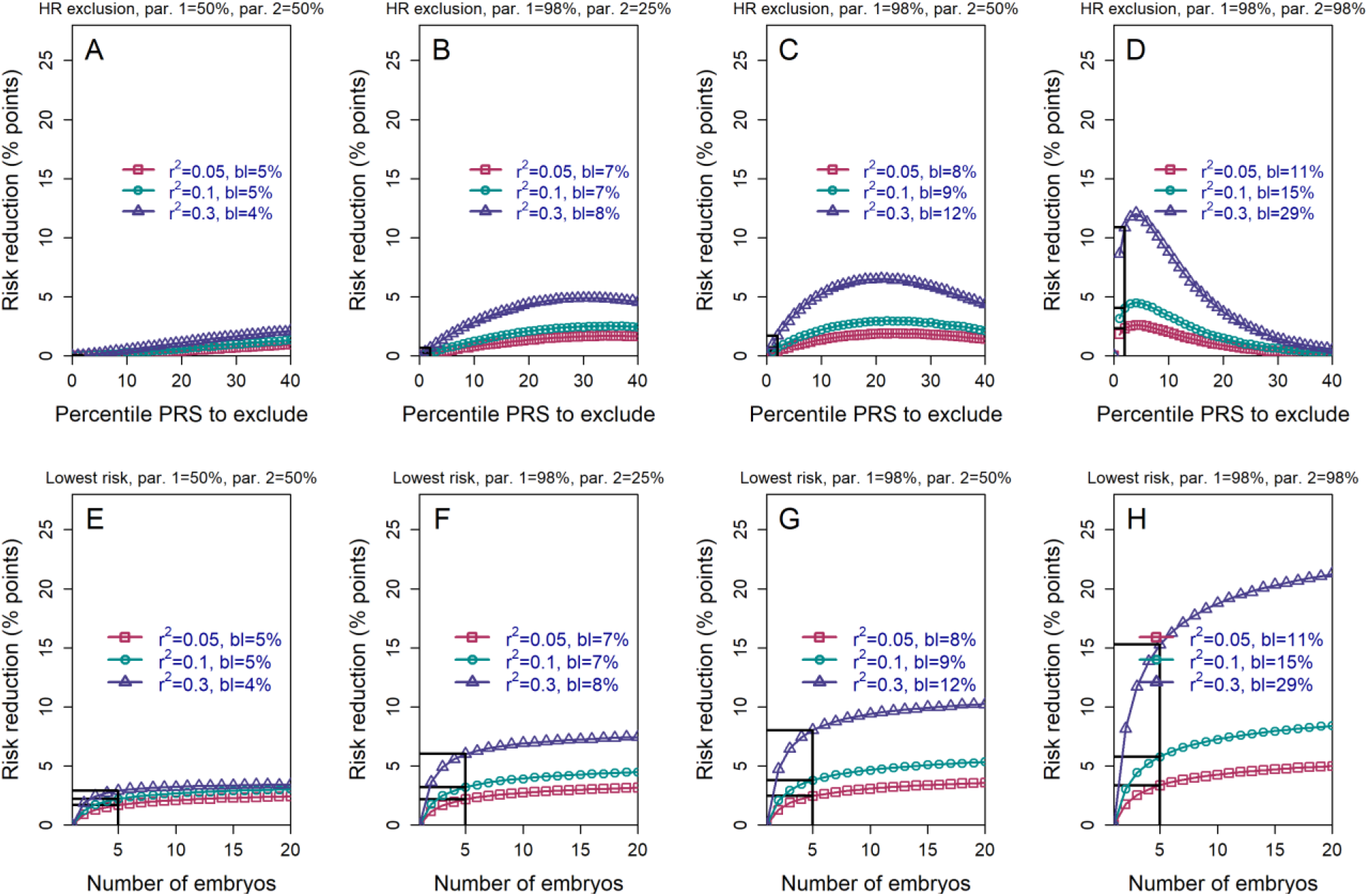
The *absolute* risk reduction when the polygenic risk scores of the parents are known. All details are the same as in Figure 3, except that the *absolute* (rather than the relative) risk reduction is shown. The absolute risk reduction is defined as the difference between the baseline disease risk (given the parental PRSs; legends) and the risk following either strategy of embryo selection. It is plotted as percentage points.

**Figure 6 – Figure Supplement 1.**
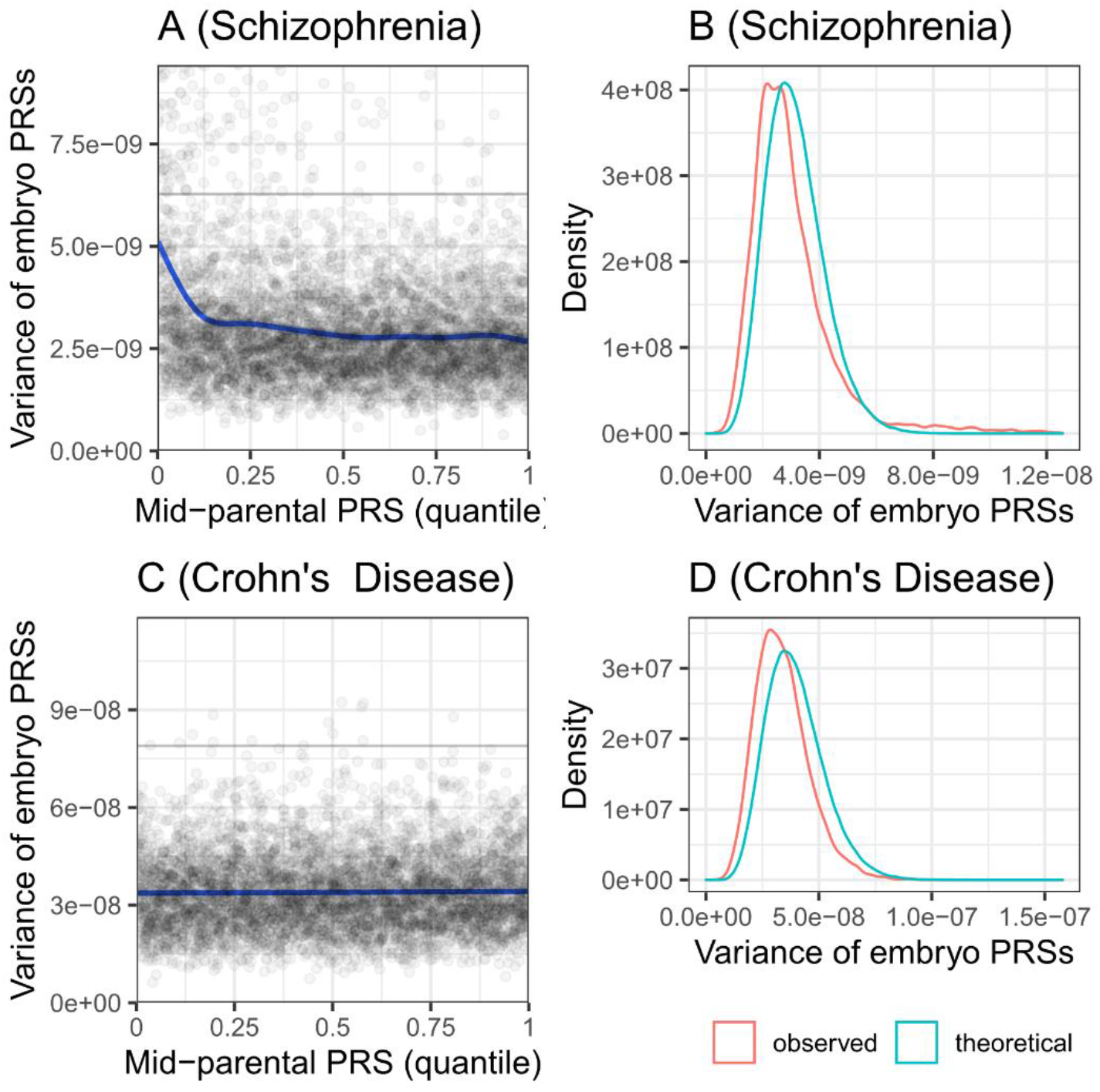
The variance of the PRS across simulated embryos. Panels (A) and (B) are for schizophrenia, while panels (C) and (D) are for Crohn’s disease. Panels (A) and (C) show the variance of the PRS across the *n* = 20 embryos of each simulated family, vs the quantile of the average parental PRS. The plots show results for 5,000 simulated families for each disease. The solid line shows smoothing cubic splines, fitted using a generalized additive model. The horizontal gray lines in (A) and (C) show the variance of the PRS in the parental population. According to the theory, the variance should be independent of the average parental PRS. Indeed, the variance is constant across average parental PRSs for Crohn’s disease. However, the variance is slightly decreasing with the average parental PRSs for schizophrenia, although the deviation is prominent only at the lowest decile. Panels (B) and (D) show the distribution of the variances across the same simulated families. The theoretical distribution is *χ*^2^ with *n* − 1 = 19 degrees of freedom, scaled by *n* − 1 and multiplied by half the variance of the PRS in the “parental” cohort. The empirical distribution (red) is very close in location and in shape to the expected distribution (cyan), although slightly shifted to lower variances.

