## Supplementary material for "Utility of polygenic embryo screening for disease depends on the selection strategy": Methods Appendix

#### 1 The liability threshold model

The liability threshold model (LTM) is a classic model in quantitative genetics (Dempster and Lerner, 1950; Falconer, 1965; Lynch and Walsh, 1998), and is also commonly used to analyze modern data (e.g., (Wray and Goddard, 2010; So et al., 2011; Lee et al., 2011, 2012; Do et al., 2012; Hayeck et al., 2017; Weissbrod et al., 2018; Hujoel et al., 2020)). Under the LTM, a disease has an underlying "liability", which is normally distributed in the population, and is the sum of two components: genetic and non-genetic (the environment). Further, the LTM assumes an "infinitesimal", or "polygenic" genetic basis, under which a very large number of genetic variants of small effect combine to form the genetic component. An individual is affected if his/her total liability (genetic + environmental) exceeds a threshold.

Mathematically, if we denote the liability as $y$, the LTM can be written as

$$y = g + \epsilon, \tag{1}$$

where $y \sim N(0,1)$ is a standard normal variable, $g \sim N(0,h^2)$ is the genetic component, with variance equal to the heritability $h^2$, and $\epsilon \sim N(0,1-h^2)$ is the non-genetic component. In practice, we cannot measure the genetic component, but only estimate it imprecisely with a polygenic risk score, denoted $s$. Following previous work (So et al., 2011; Do et al., 2012; Lee et al., 2012; Treff et al., 2019a; Karavani et al., 2019), we assume that the LTM can be written, similarly to Eq. (1), as

$$y = s + e, \tag{2}$$

where $y \sim N(0,1)$ as above, $s \sim N(0,r_{\text{ps}}^2)$, where $r_{\text{ps}}^2$ is the proportion of the variance in liability explained by the score, and $e \sim N(0,1-r_{\text{ps}}^2)$ is the residual of the regression of the liability on $s$ (and is uncorrelated with $s$), representing environmental effects *as well as* genetic factors not accounted for by the score.

An individual is affected whenever his/her liability exceeds a threshold. The threshold is selected such that the proportion of affected individuals is equal to the prevalence $K$, i.e., it is equal to $z_K$, the $(1-K)$-quantile of a standard normal variable. Thus,

$$P(\text{disease}) = P(y > z_K) = K. \tag{3}$$

The model is illustrated in Figure 1B of the main text.

### 2 A model for the scores of $n$ IVF embryos

Consider the polygenic risk scores (for a disease of interest) of $n$ IVF embryos of given parents. We assume no information is known about the parents, or, in other words, that the parents are randomly and independently drawn from the population. The scores of the embryos have a multivariate normal distribution,

$$\boldsymbol{s} = (s_1, \ldots, s_n) = \text{MVN}(\mathbf{0}_n, \boldsymbol{\Sigma}), \tag{4}$$

where the means form a vector $\mathbf{0}_n$ of $n$ zeros, and the $n \times n$ covariance matrix is

$$\boldsymbol{\Sigma} = r_{\text{ps}}^2 \begin{pmatrix} 1 & \frac{1}{2} & \cdots & \frac{1}{2} \\ \frac{1}{2} & 1 & \cdots & \frac{1}{2} \\ \cdots & \cdots & \cdots & \cdots \\ \frac{1}{2} & \frac{1}{2} & \cdots & 1 \end{pmatrix}. \tag{5}$$

The diagonal elements of the matrix are simply the variances of the individual scores of each embryo. The off-diagonal elements represent the covariance between the scores of the embryos, who are genetically siblings. Based on standard quantitative genetic theory (Lynch and Walsh, 1998) (see also our previous paper (Karavani et al., 2019)), the covariance between the scores of two siblings is $\text{Cov}(s_i, s_j) = \frac{1}{2}\text{Var}(s)$, and hence the off-diagonal elements follow. [The non-score components (the $e$ terms in Eq. (2)) are also correlated. The correlation between the genetic components of $e$ is modeled in Section 6. Modeling the correlation between the environmental components was unnecessary in this paper – see Section 10.]

As we showed in our previous work (Karavani et al., 2019), the scores can be written as a sum of two independent multivariate normal variables, $\boldsymbol{s} = \boldsymbol{x} + \boldsymbol{c}$, with

$$\boldsymbol{x} = (x_1, \ldots, x_n) \sim \text{MVN}\left(\mathbf{0}_n, \frac{r_{\text{ps}}^2}{2}\boldsymbol{I}_n\right) \text{ and}$$
$$\boldsymbol{c} = (c_1, \ldots, c_n) \sim \text{MVN}\left(\mathbf{0}_n, \frac{r_{\text{ps}}^2}{2}\boldsymbol{J}_n\right), \tag{6}$$

where $\mathbf{0}_n$ is a vector of zeros of length $n$, $\boldsymbol{I}_n$ is the $n \times n$ identity matrix, and $\boldsymbol{J}_n$ is the $n \times n$ matrix of all ones. The $x_i$'s and $c_i$'s have the same marginal distribution, namely normal with mean zero and variance $r_{\text{ps}}^2/2$ each. However, the $x_i$'s are independent, whereas $\boldsymbol{c}$ has a constant covariance matrix, which means that the $c_i$'s are $n$ identical copies of the same random variable,

$$c_1 \sim N\left(0, \frac{r_{\text{ps}}^2}{2}\right) \text{ and } c_2 = c_3 = \cdots = c_n = c_1 \equiv c. \tag{7}$$

Thus, for each embryo $i = 1, \dots, n$,

$$s_i = x_i + c. \tag{8}$$

#### 2.1 An alternative interpretation: conditioning on the average parental scores

The decomposition of the score in Eq. (8) can also be interpreted as conditioning on the average score of the parents. To see that, write the maternal score as $s_m$ and the paternal score as $s_f$. The variables $(s_i, s_m, s_f)$ have a multivariate normal distribution,

$$(s_i, s_m, s_f) \sim \text{MVN}\left(\begin{pmatrix}0\\0\\0\end{pmatrix}, \begin{pmatrix} r_{\text{ps}}^2 & \frac{r_{\text{ps}}^2}{2} & \frac{r_{\text{ps}}^2}{2} \\ \frac{r_{\text{ps}}^2}{2} & r_{\text{ps}}^2 & 0 \\ \frac{r_{\text{ps}}^2}{2} & 0 & r_{\text{ps}}^2 \end{pmatrix}\right). \tag{9}$$

In the above equation, the variances of all scores are equal to $r_{\text{ps}}^2$. The covariance terms are $\text{Cov}\,(s_i, s_m) = \text{Cov}\,(s_i, s_f) = \frac{1}{2}\text{Var}\,(s) = \frac{r_{\text{ps}}^2}{2}$, as the relatedness between between parent and child is the same as for a pair of siblings. We assume no correlation between the scores of the parents (i.e., no assortative mating, see Section 10 for discussion). We are now interested in the conditional density of $s_i$ given $s_m$ and $s_f$. Using standard results for multivariate normal distributions, the conditional density of $s_i$ is $N(\mu, \sigma^2)$, where,

$$\mu = \boldsymbol{\Sigma}_{12}\boldsymbol{\Sigma}_{22}^{-1}\begin{pmatrix}s_m\\s_f\end{pmatrix},$$
$$\sigma^2 = \boldsymbol{\Sigma}_{11} - \boldsymbol{\Sigma}_{12}\boldsymbol{\Sigma}_{22}^{-1}\boldsymbol{\Sigma}_{21}, \tag{10}$$

and

$$\boldsymbol{\Sigma}_{11} = r_{\text{ps}}^2, \boldsymbol{\Sigma}_{12} = \begin{pmatrix}\frac{r_{\text{ps}}^2}{2} & \frac{r_{\text{ps}}^2}{2}\end{pmatrix}, \boldsymbol{\Sigma}_{21} = \begin{pmatrix}\frac{r_{\text{ps}}^2}{2}\\ \frac{r_{\text{ps}}^2}{2}\end{pmatrix}, \boldsymbol{\Sigma}_{22} = \begin{pmatrix} r_{\text{ps}}^2 & 0 \\ 0 & r_{\text{ps}}^2\end{pmatrix}. \tag{11}$$

These matrices are the blocks forming the covariance matrix in Eq. (9). Carrying out the matrix calculations, we obtain

$$\mu = \frac{s_m + s_f}{2},$$
$$\sigma^2 = \frac{r_{\text{ps}}^2}{2}. \tag{12}$$

Thus,

$$s_i \sim N\left(\frac{s_m + s_f}{2}, \frac{r_{\text{ps}}^2}{2}\right) \equiv N(c, r_{\text{ps}}^2/2), \tag{13}$$

where we defined the shared component $c \equiv \frac{s_m+s_f}{2}$ as the average parental score. The variance of $c$ itself, across the population, is $\text{Var}\left(\frac{s_m+s_f}{2}\right) = \frac{2\text{Var}(s)}{4} = r_{\text{ps}}^2/2$. Thus, $c \sim N(0, r_{\text{ps}}^2/2)$. In a given family, $c$ is the same across all embryos. Thus, Eq. (13) is equivalent to $s_i = c + x_i$, with $c \sim N(0, r_{\text{ps}}^2/2)$ and $x_i \sim N(0, r_{\text{ps}}^2/2)$ is an embryo-specific component.

An analogous result holds for the total genetic component of the embryo, $g_i$, simply by replacing the proportion of variance explained by the score ($r_{\text{ps}}^2$) with the heritability ($h^2$). In other words, if $g_m$ and $g_f$ are the maternal and paternal genetic components, respectively, then

$$g_i \sim N\left(\frac{g_m + g_f}{2}, \frac{h^2}{2}\right). \tag{14}$$

### 3 The disease risk when implanting the embryo with the lowest risk

We assume next that we select for implantation the embryo with the lowest polygenic risk score for the disease of interest. Our goal will be to calculate the probability of that embryo to be affected. Since $s_i = x_i + c$, the score of the selected embryo satisfies

$$\begin{aligned} s_{\min} &= \min(x_1 + c, \dots, x_n + c) \\ &= \min(x_1, \dots, x_n) + c \\ &= x_{\min} + c, \end{aligned} \tag{15}$$

where we defined $x_{\min} = \min(x_1, \dots, x_n)$. Denote by $i^*$ the index of the selected embryo ($x_{i^*} = x_{\min}$). The liability of the embryo with the lowest risk is thus

$$\begin{aligned} y_{i^*} &= s_{\min} + e_{i^*} \\ &= x_{\min} + c + e_{i^*} \\ &= x_{\min} + \tilde{e}, \end{aligned} \tag{16}$$

where $e_i$ is the non-score component of embryo $i$, and $\tilde{e} = c + e_{i^*}$. We have,

$$\text{Var}(\tilde{e}) = \text{Var}(c) + \text{Var}(e_{i^*}) = \frac{r_{\text{ps}}^2}{2} + (1 - r_{\text{ps}}^2) = 1 - \frac{r_{\text{ps}}^2}{2}. \tag{17}$$

Therefore, the liability of the selected embryo can be written as a sum of two (independent) variables: $x_{\min}$, which is the minimum of $n$ *independent* (zero mean) normal variables with variance $r_{\text{ps}}^2/2$ each; and $\tilde{e}$, which is a normal variable with (zero mean and) variance $1 - r_{\text{ps}}^2/2$.

The distribution of $x_{\min}$ can be computed based on the theory of order statistics,

$$P(x_{\min} > t) = [P(x > t)]^n = \left[1 - \Phi\left(\frac{t}{r_{\text{ps}}/\sqrt{2}}\right)\right]^n. \tag{18}$$

In the above equation, the minimum of $n$ variables is greater than $t$ if and only if all variables are greater than $t$. The distribution of each $x$ is normal with zero mean and variance $r_{\text{ps}}^2/2$, and hence $P(x > t) = 1 - \Phi\left(\frac{t}{r_{\text{ps}}/\sqrt{2}}\right)$, where $\Phi(\cdot)$ is the cumulative probability distribution (CDF) of a standard normal variable.

We can now compute the probability of the selected embryo to be affected by demanding that the total liability is greater than the threshold $z_K$. Denote the probability of disease as $P_s(\text{disease})$ ($s$ stands for *s*elected). Conditional on $\tilde{e}$,

$$\begin{aligned} P_s(\text{disease} \mid \tilde{e}) &= P(y_{i^*} > z_K \mid \tilde{e}) \\ &= P(x_{\min} + \tilde{e} > z_K) \\ &= P(x_{\min} > z_K - \tilde{e}) \\ &= \left[1 - \Phi\left(\frac{z_K - \tilde{e}}{r_{\text{ps}}/\sqrt{2}}\right)\right]^n, \end{aligned} \tag{19}$$

where in the fourth line, we used Eq. (18). Next, denote by $f(\tilde{e})$ the density of $\tilde{e}$, and by $\phi(\cdot)$ the probability density function of a standard normal variable. Given that $\tilde{e} \sim N(0, 1 - r_{\text{ps}}^2/2)$,

$$\begin{aligned} P_s(\text{disease}) &= \int_{-\infty}^{\infty} P_s(\text{disease} \mid \tilde{e}) f(\tilde{e}) d\tilde{e} \\ &= \int_{-\infty}^{\infty} \left[1 - \Phi\left(\frac{z_K - \tilde{e}}{r_{\text{ps}}/\sqrt{2}}\right)\right]^n \frac{1}{\sqrt{1 - r_{\text{ps}}^2/2}} \phi\left(\frac{\tilde{e}}{\sqrt{1 - r_{\text{ps}}^2/2}}\right) d\tilde{e} \\ &= \int_{-\infty}^{\infty} \left[1 - \Phi\left(\frac{z_K - t\sqrt{1 - r_{\text{ps}}^2/2}}{r_{\text{ps}}/\sqrt{2}}\right)\right]^n \phi(t) dt. \end{aligned} \tag{20}$$

In the third line, we changed variables: $t = \tilde{e}/\sqrt{1 - r_{\text{ps}}^2/2}$. Eq. (20) is our final expression for the probability of the embryo with the lowest score to be affected.

#### 3.1 The risk reduction when conditioning on the mean parental score

Consider the case when $c$ is given, or, in other words, when we know the mean parental polygenic score. Let us compute the disease risk in such a case. We start from Eq. (16),

$$\begin{aligned} y_{i^*} &= s_{\min} + e_{i^*} \\ &= x_{\min} + c + e_{i^*}. \end{aligned} \tag{21}$$

Then,

$$\begin{aligned} P_s(\text{disease} \,|\, c, e_{i^*}) &= P(y_{i^*} > z_K \,|\, e_{i^*}) \\ &= P(x_{\min} + c + e_{i^*} > z_K) \\ &= P(x_{\min} > z_K - c - e_{i^*}) \\ &= \left[1 - \Phi\left(\frac{z_K - c - e_{i^*}}{r_{\text{ps}}/\sqrt{2}}\right)\right]^n, \end{aligned} \tag{22}$$

where in the last line, we used Eq. (18).

Finally, with $f(e_{i^*})$ denoting the density of $e_{i^*}$, and recalling that $e_{i^*} \sim N\left(0, 1 - r_{\text{ps}}^2\right)$,

$$\begin{aligned} P_s(\text{disease} \,|\, c) &= \int_{-\infty}^{\infty} P_s(\text{disease} \,|\, c, e_{i^*}) f(e_{i^*}) de_{i^*} \\ &= \int_{-\infty}^{\infty} \left[1 - \Phi\left(\frac{z_K - c - e_{i^*}}{r_{\text{ps}}/\sqrt{2}}\right)\right]^n \frac{1}{\sqrt{1 - r_{\text{ps}}^2}} \phi\left(\frac{e_{i^*}}{\sqrt{1 - r_{\text{ps}}^2}}\right) de_{i^*} \\ &= \int_{-\infty}^{\infty} \left[1 - \Phi\left(\frac{z_K - c - t\sqrt{1 - r_{\text{ps}}^2}}{r_{\text{ps}}/\sqrt{2}}\right)\right]^n \phi(t) dt, \end{aligned} \tag{23}$$

where in the last line, we changed variables, $t = e_{i^*}/\sqrt{1 - r_{\text{ps}}^2}$. Eq. (23) thus provides the probability of disease when we are given the mean parental score $c$.

### 4 The disease risk when excluding high-risk embryos

We now consider the selection strategy in which the implanted embryo is selected at random, as long as its risk score is not particularly high. Specifically, we assume that whenever possible, embryos at the top $q$ risk percentiles are excluded. When *all* embryos have high risk, we assume that a random embryo is selected. Let $z_q$ be the $(1 - q)$-quantile of the standard normal distribution. The variance of the score is $r_{\text{ps}}^2$, and therefore, the score of the selected embryo must be lower than $z_q r_{\text{ps}}$.

To compute the disease risk in this case, we first condition on the shared, family-specific component $c$. We later integrate over $c$ to derive the risk across the population. Denote by $x_s$ the value of $x$ for the *selected* embryo, and for the moment, also condition on $x_s$. We have,

$$\begin{aligned}
P_s(\text{disease} \,|\, x_s, c) &= P(y > z_K \,|\, c) \\
&= P(s + e > z_K \,|\, c) \\
&= P(x_s + c + e > z_K) \\
&= P(e > z_K - x_s - c) \\
&= 1 - \Phi\left(\frac{z_K - x_s - c}{\sqrt{1 - r_{\text{ps}}^2}}\right),
\end{aligned} \tag{24}$$

To obtain $P_s(\text{disease} \,|\, c)$, we need to integrate over $f(x_s)$, the density of $x_s$. In fact, $f(x_s)$ is a mixture of two distributions, depending on whether or not all embryos were high risk. Denote by $H$ the event that all embryos are high risk, and let us first compute the probability of $H$. Recall that given $c$, the scores of all embryos, $s_i = x_i + c$, are independent. The event $H$ is equivalent to the intersection of the *independent* events $\{s_i > z_q r_{\text{ps}}\}$ for $i = 1, \dots, n$. Thus, recalling that $x_i \sim N(0, r_{\text{ps}}^2/2)$,

$$\begin{aligned}
P(H) &= \prod_{i=1}^{n} P(s_i > z_q r_{\text{ps}}) \\
&= \prod_{i=1}^{n} P(x_i + c > z_q r_{\text{ps}}) \\
&= \prod_{i=1}^{n} P(x_i > z_q r_{\text{ps}} - c) \\
&= \left[1 - \Phi\left(\frac{z_q r_{\text{ps}} - c}{r_{\text{ps}}/\sqrt{2}}\right)\right]^n .
\end{aligned} \tag{25}$$

Given $H$, we know that all scores were higher than the cutoff, i.e., that $x_i > z_q r_{\text{ps}} - c$ for all $i = 1, \dots, n$. An embryo is then selected at random. Thus, $x_s$, the value of $x$ of the selected embryo, is a realization of a normal random variable truncated from below. Specifically, if $f_x(\cdot)$ is the unconditional density of $x$, then for $x_s > z_q r_{\text{ps}} - c$,

$$f(x_s \,|\, H) = \frac{f_x(x_s)}{P(x > z_q r_{\text{ps}} - c)} = \frac{\frac{1}{r_{\text{ps}}/\sqrt{2}} \phi\left(\frac{x_s}{r_{\text{ps}}/\sqrt{2}}\right)}{1 - \Phi\left(\frac{z_q r_{\text{ps}} - c}{r_{\text{ps}}/\sqrt{2}}\right)}. \tag{26}$$

In the case $H$ did not occur, we select an embryo at random among embryos with score $s_i < z_q r_{\text{ps}}$, i.e., $x_i < z_q r_{\text{ps}} - c$. The density of $x_s$ is again, analogously to the above case, a realization of a normal random variable, but this time truncated from above. For $x_s < z_q r_{\text{ps}} - c$,

$$f(x_s \,|\, \overline{H}) = \frac{f_x(x_s)}{P(x < z_q r_{\text{ps}} - c)} = \frac{\frac{1}{r_{\text{ps}}/\sqrt{2}} \phi\left(\frac{x_s}{r_{\text{ps}}/\sqrt{2}}\right)}{\Phi\left(\frac{z_q r_{\text{ps}} - c}{r_{\text{ps}}/\sqrt{2}}\right)}. \tag{27}$$

Using these results, we can write the density of $x_s$ when conditioning only on $c$,

$$\begin{aligned} f(x_s) &= \begin{cases} f(x_s \,|\, H)P(H) + 0 \cdot P(\overline{H}) & \text{for } x_s > z_q r_{\text{ps}} - c \\ 0 \cdot P(H) + f(x_s \,|\, \overline{H})P(\overline{H}) & \text{for } x_s < z_q r_{\text{ps}} - c \end{cases} \\ &= \begin{cases} \frac{\frac{1}{r_{\text{ps}}/\sqrt{2}} \phi\left(\frac{x_s}{r_{\text{ps}}/\sqrt{2}}\right)}{1 - \Phi\left(\frac{z_q r_{\text{ps}} - c}{r_{\text{ps}}/\sqrt{2}}\right)} \left[1 - \Phi\left(\frac{z_q r_{\text{ps}} - c}{r_{\text{ps}}/\sqrt{2}}\right)\right]^n & \text{for } x_s > z_q r_{\text{ps}} - c \\ \frac{\frac{1}{r_{\text{ps}}/\sqrt{2}} \phi\left(\frac{x_s}{r_{\text{ps}}/\sqrt{2}}\right)}{\Phi\left(\frac{z_q r_{\text{ps}} - c}{r_{\text{ps}}/\sqrt{2}}\right)} \left\{1 - \left[1 - \Phi\left(\frac{z_q r_{\text{ps}} - c}{r_{\text{ps}}/\sqrt{2}}\right)\right]^n\right\} & \text{for } x_s < z_q r_{\text{ps}} - c \end{cases} \end{aligned} \tag{28}$$

We can now integrate over all $x_s$, still conditioning on $c$, and using Eq. (24) and some algebra,

$$\begin{aligned} &P_s(\text{disease} \,|\, c) = \int_{-\infty}^{\infty} f(x_s) P_s(\text{disease} \,|\, x_s, c) dx_s \\ &= \int_{-\infty}^{z_q r_{\text{ps}} - c} \frac{\frac{1}{r_{\text{ps}}/\sqrt{2}} \phi\left(\frac{x_s}{r_{\text{ps}}/\sqrt{2}}\right)}{\Phi\left(\frac{z_q r_{\text{ps}} - c}{r_{\text{ps}}/\sqrt{2}}\right)} \left\{1 - \left[1 - \Phi\left(\frac{z_q r_{\text{ps}} - c}{r_{\text{ps}}/\sqrt{2}}\right)\right]^n\right\} \left[1 - \Phi\left(\frac{z_K - x_s - c}{\sqrt{1 - r_{\text{ps}}^2}}\right)\right] dx_s \\ &+ \int_{z_q r_{\text{ps}} - c}^{\infty} \frac{\frac{1}{r_{\text{ps}}/\sqrt{2}} \phi\left(\frac{x_s}{r_{\text{ps}}/\sqrt{2}}\right)}{1 - \Phi\left(\frac{z_q r_{\text{ps}} - c}{r_{\text{ps}}/\sqrt{2}}\right)} \left[1 - \Phi\left(\frac{z_q r_{\text{ps}} - c}{r_{\text{ps}}/\sqrt{2}}\right)\right]^n \left[1 - \Phi\left(\frac{z_K - x_s - c}{\sqrt{1 - r_{\text{ps}}^2}}\right)\right] dx_s \\ &= \int_{-\infty}^{\infty} \eta(t, \gamma(c)) \xi(t, c) dt, \end{aligned} \tag{29}$$

where we defined

$$\begin{aligned} \xi(t, c) &= \phi(t) \left[1 - \Phi\left(\frac{z_K - t r_{\text{ps}}/\sqrt{2} - c}{\sqrt{1 - r_{\text{ps}}^2}}\right)\right], \\ \eta(t, \gamma) &= \begin{cases} \frac{1 - [1 - \Phi(\gamma)]^n}{\Phi(\gamma)} & \text{for } t < \gamma, \\ [1 - \Phi(\gamma)]^{n-1} & \text{for } t > \gamma. \end{cases}, \text{ and} \\ \gamma(c) &= \sqrt{2} z_q - \frac{c}{r_{\text{ps}}/\sqrt{2}}. \end{aligned} \tag{30}$$

Eq. (29) provides an expression for the probability of a disease given the mean parental score $c$.

Finally, we can integrate over all $c$ in order to obtain the probability of disease in the population. Recalling that $c \sim N(0, r_{\text{ps}}^2/2)$ and denoting its density as $f(c)$, and again after some algebra,

$$\begin{aligned} P_s(\text{disease}) &= \int_{-\infty}^{\infty} P_s(\text{disease} \mid c) f(c) dc \\ &= \int_{-\infty}^{\infty} \phi(u) \left[ \int_{-\infty}^{\infty} \eta(t, \beta(u)) \zeta(u,t) dt \right] du, \end{aligned} \tag{31}$$

where we defined

$$\begin{aligned} \zeta(u,t) &= \phi(t) \left[ 1 - \Phi\left( \frac{z_K - (u+t) r_{\text{ps}}/\sqrt{2}}{\sqrt{1 - r_{\text{ps}}^2}} \right) \right], \\ \beta(u) &= \sqrt{2} z_q - u, \end{aligned} \tag{32}$$

and $\eta(t, \cdot)$ was defined in Eq. (30) above. Eq. (31) is our final expression for the probability of an embryo to be affected after being selected randomly among non-high-risk embryos.

### 5 The relative risk reduction

We define the relative risk reduction (RRR) as follows. We are given the prevalence $K$ and the probability of the selected embryo to be affected $P_s(\text{disease})$ (averaged over the population). Then,

$$\text{RRR} = \frac{K - P_s(\text{disease})}{K} = 1 - \frac{P_s(\text{disease})}{K}. \tag{33}$$

The absolute risk reduction (ARR) is similarly defined as $K - P_s(\text{disease})$. For example, if a disease has prevalence of 5% and an embryo selected based on PRS has an average probability of 3% to be affected, the relative risk reduction is 40%, while the absolute risk reduction is 2%.

To use Eq. (33), $P_s(\text{disease})$ is given by Eq. (20) for the *lowest-risk prioritization* strategy, and by Eq. (31) for the *high-risk exclusion* strategy. We solve the integrals in these equations numerically in R using the function `integrate` (see Section 11).

#### 5.1 The *per-couple* relative risk reduction

The RRR, as defined in Eq. (33), is the (complement of the) ratio between two *average* risks: the average risk of a random couple that would select an embryo based on its PRS, and the average risk of a random couple that would select an embryo at random. It can also be seen as the relative risk reduction between the risks in two hypothetical "populations": one in which all embryos are selected based on a PRS-based strategy, and one in which all embryos are selected at random.

However, a shortcoming of the population-level RRR definition is that it does not provide information on the risk reduction expected for *individual couples*. In other words, a given couple may wish to know the extent to which they can reduce disease risk in their children by electing to select an embryo based on PRS. Conveniently, the only relevant information that characterizes the potential risk reduction for a given couple is $c$, the average parental score.

We define the *per-couple* relative risk reduction, or pcRRR($c$), as

$$\text{pcRRR}(c) = \frac{P_r(\text{disease}\,|\,c) - P_s(\text{disease}\,|\,c)}{P_r(\text{disease}\,|\,c)} = 1 - \frac{P_s(\text{disease}\,|\,c)}{P_r(\text{disease}\,|\,c)}, \tag{34}$$

where $P_r(\text{disease}\,|\,c)$ is the "baseline" risk, i.e., the probability of disease of a random embryo ($r$ stands for $r$andom; this can also be seen as the risk in natural procreation). Note that we can similarly define the absolute risk reduction (ARR) as $P_r(\text{disease}\,|\,c) - P_s(\text{disease}\,|\,c)$.

We have already computed $P_s(\text{disease}\,|\,c)$ for the two selection strategies (Eqs. (23) and (29)). To compute $P_r(\text{disease}\,|\,c)$, we write the liability of a random embryo as

$$\begin{aligned} y &= s + e \\ &= x + c + e \\ &= \tilde{x} + c, \end{aligned} \tag{35}$$

where we defined $\tilde{x} = x + e$. $\text{Var}(\tilde{x}) = \text{Var}(x) + \text{Var}(e) = r_{\text{ps}}^2/2 + 1 - r_{\text{ps}}^2 = 1 - r_{\text{ps}}^2/2$, and thus, $\tilde{x} \sim N\left(0, 1 - r_{\text{ps}}^2/2\right)$. The conditional probability of disease is

$$\begin{aligned} P_r(\text{disease}\,|\,c) &= P(y > z_K\,|\,c) \\ &= P(\tilde{x} + c > z_K) \\ &= P(\tilde{x} > z_K - c) \\ &= 1 - \Phi\left(\frac{z_K - c}{\sqrt{1 - r_{\text{ps}}^2/2}}\right). \end{aligned} \tag{36}$$

#### 5.2 The distribution of the *per-couple* relative risk reduction

We can compute the probability density of pcRRR($c$) across all couples in the population, $f_{pc}(x)$, as follows,

$$f_{pc}(x) = \int_{-\infty}^{\infty} \delta\left(x - \text{pcRRR}(c)\right) f(c) dc, \tag{37}$$

where $\delta(x)$ is Dirac's delta function, $c$ is the parental average score, and $f(c) \sim N\left(0, r_{\text{ps}}^2/2\right)$ is its density. For computing $f_{pc}(x)$ numerically, we sum over $10^4$

quantiles of $c$ (which by definition have equal probability), and then compute the probability of the pcRRR to have value within each bin,

$$P(\text{pcRRR} \in [r_1, r_2]) = \frac{1}{10^4} \sum_{i=1}^{10^4} \mathbf{1}_{\text{pcRRR}(c_i) \in [r_1, r_2]}, \tag{38}$$

where $\mathbf{1}$ is the indicator variable, and $c_i$ is the $i/10^4$ quantile of $c$ (a value such that $c$ is less than $c_i$ with probability $(i - 0.5)/10^4$).

The average pcRRR across all couples is

$$\langle \text{pcRRR} \rangle = \int_{-\infty}^{\infty} \text{pcRRR}(c) f(c) dc. \tag{39}$$

Numerically,

$$\langle \text{pcRRR} \rangle = \frac{1}{10^4} \sum_{i=1}^{10^4} \text{pcRRR}(c_i). \tag{40}$$

Note that Eq. (39) is an average of ratios. This is in contrast to Eq. (33), which a ratio of averages. As such, those average risk reductions are not expected to be identical. Empirically, given that pcRRR($c$) depends only weakly on $c$, we found that differences were small. For example, $\langle$pcRRR$\rangle$ was higher than the RRR from Eq. (33) by $\approx 0.01$ for $r_{\text{ps}}^2 \leq 0.1$ (for $K = 0.01, 0.05, 0.2$); when $K = 0.05$ and $r_{\text{ps}}^2 = 0.1$, $\langle$pcRRR$\rangle$ was 0.48, while the RRR was 0.47. Differences were larger for $r_{\text{ps}}^2 = 0.3$; for example, for $K = 0.05$, $\langle$pcRRR$\rangle$ was 0.77, while the RRR was 0.72.

#### 5.3 The *per-batch* relative risk reduction

The pcRRR, i.e., Eq. (34), can be interpreted as follows. A given couple can choose between two options: either generate embryos by IVF and select an embryo based on its PRS, or select an embryo at random (=conceive naturally). The pcRRR quantifies the risk reduction between the outcomes under these two choices. For each choice, the risk is computed by averaging over all possible embryos that may have been generated in an IVF cycle. However, one may also wish to quantify the variability of the outcome for a given couple. This could be accomplished as follows: for each couple *and* for each batch of $n$ embryos, compute the relative risk reduction when selecting an embryo based on PRS vs when selecting at random. We define this quantity as the *per-batch* relative risk reduction, or pbRRR.

Modeling the pbRRR is straightforward using our framework. Given the scores of the embryos, $s_1, \ldots, s_n$, the selected embryo is immediately determined for the *lowest-risk prioritization* strategy. For the *high-risk exclusion* strategy, the selected embryo can be, with equal probability, any of the embryos that are not high risk (or any embryo if all embryos are high risk). For random selection, the selected embryo can be any embryo with equal probability. Given the score of the selected embryo, $s_{i^*}$, and given the non-score component, $e \sim N(0, 1 - r_{\text{ps}}^2)$, the probability of disease of the selected embryo is

$$\begin{aligned} P(\text{disease}) &= P(y > z_K \mid s_{i^*}) \\ &= P(s_{i^*} + e > z_K) \\ &= P(e > z_K - s_{i^*}) \\ &= 1 - \Phi\left(\frac{z_K - s_{i^*}}{\sqrt{1 - r_{\text{ps}}^2}}\right). \end{aligned} \quad (41)$$

The probability density of the scores is then given by Eq. (13). The distribution of the pbRRR across batches of embryos can then be computed by integrating over all possible sets of $n$ scores, similarly to Eq. (37). However, this would be tedious in practice, and we do not pursue this direction here.

### 6 The risk reduction conditional on family history

In the following, we compute the relative risk reduction when the disease status of the parents is given.

#### 6.1 Model

Let us rewrite our model for the liability as

$$y = s + w + \epsilon. \quad (42)$$

Here, $w$ represents all genetic factors not included in the score. We keep track of both $s$ and $w$, because both are inherited, and hence, information on the disease status of the parents will be informative on their values in children (see below). However, we need to track each term separately because selection is only based on $s$. As in Section 1, we assume $s$, $w$, and $\epsilon$ are independent, $y \sim N(0, 1)$, $s \sim N(0, r_{\text{ps}}^2)$, and $\epsilon \sim N(0, 1 - h^2)$, and thus $w \sim N(0, h^2 - r_{\text{ps}}^2)$.

We derive the risk to the embryos in two main steps. First, we assume that the values of $s$ and $w$ are known for each parent, and compute the risk of the embryo under each selection strategy (*lowest-risk prioritization*, *high-risk exclusion*, or random selection). Then, we derive the posterior distribution of the parental genetic components given the parental disease status, and integrate over these components to obtain the final risk estimate.

#### 6.2 The risk of the selected embryo given its score

Denote the maternal score as $s_m$ and the paternal score as $s_f$, denote similarly $w_m$ and $w_f$, and assume that they are given. Also denote $g_m = s_m + w_m$ and

$g_f = s_f + w_f$. As we explained in Section 2.1, for any child $i$, the distribution of the score $s_i$ is

$$s_i \sim N\left(\frac{s_m + s_f}{2}, \frac{r_{\text{ps}}^2}{2}\right) \text{ or } s_i = c + x_i, \tag{43}$$

where $c = (s_m + s_f)/2$ and $x_i \sim N(0, r_{\text{ps}}^2/2)$. Similarly, the distribution of the non-score genetic component is

$$w_i \sim N\left(\frac{w_m + w_f}{2}, \frac{h^2 - r_{\text{ps}}^2}{2}\right) \text{ or } w_i = \frac{w_m + w_f}{2} + v_i, \tag{44}$$

where $v_i \sim N\left(0, (h^2 - r_{\text{ps}}^2)/2\right)$.

Given the parental genetic components, we can write the liability of each embryo as, for $i = 1, \ldots, n$,

$$y_i = \frac{s_m + s_f}{2} + x_i + \frac{w_m + w_f}{2} + v_i + \epsilon_i, \tag{45}$$

where $\epsilon_i \sim N(0, 1 - h^2)$. All the three random variables in the above equation ($x_i$, $v_i$, and $\epsilon_i$) are independent, and $x_i$ and $v_i$ are each independent across embryos. (It is not necessary to specify whether the $\epsilon_i$ are independent.) Denote the event that embryo $i$ is affected as $D_i$, and condition on the value of $x_i$ for that embryo. The probability of disease is

$$\begin{aligned}
P\left(D_i \,|\, s_m, w_m, s_f, w_f, x_i\right) &= P(y_i > z_K \,|\, s_m, w_m, s_f, w_f, x_i) \\
&= P\left(\frac{s_m + s_f}{2} + x_i + \frac{w_m + w_f}{2} + v_i + \epsilon_i > z_K\right) \\
&= P\left(v_i + \epsilon_i > z_K - \frac{s_m + s_f}{2} - \frac{w_m + w_f}{2} - x_i\right) \\
&= 1 - \Phi\left(\frac{z_K - \frac{s_m + s_f}{2} - \frac{w_m + w_f}{2} - x_i}{\sqrt{1 - h^2/2 - r_{\text{ps}}^2/2}}\right).
\end{aligned} \tag{46}$$

The last line holds because $\text{Var}\left(v_i + \epsilon_i\right) = (h^2 - r_{\text{ps}}^2)/2 + (1 - h^2) = 1 - h^2/2 - r_{\text{ps}}^2/2$.

We henceforth denote $D_s$ as the event that the selected embryo is affected. In the next three subsections, we integrate the probability of the disease over $x_i$, where the distribution of $x_i$ will vary depending on the selection strategy. This will give us the disease risk given the parental genetic components.

#### 6.3 Selecting the lowest-risk embryo

Denote by $x_{i^*}$ the embryo-specific component of the embryo with the lowest such component. Recall that for each embryo, $x_i \sim N(0, r_{\text{ps}}^2/2)$. We can use the theory of order statistics, as in previous sections, to compute the density of $x_{i^*}$.

$$f(x_{i^*}) = \frac{n}{r_{\text{ps}}/\sqrt{2}}\phi\left(\frac{x_{i^*}}{r_{\text{ps}}/\sqrt{2}}\right)\left[1-\Phi\left(\frac{x_{i^*}}{r_{\text{ps}}/\sqrt{2}}\right)\right]^{n-1}. \tag{47}$$

Eq. (46) can now be integrated over all $x_{i*}$. After changing variables $t = x_{i^*}/(r_{\text{ps}}/\sqrt{2})$, we obtain

$$\begin{aligned} &P\left(D_s \,|\, s_m, w_m, s_f, w_f\right) = \\ &= \int_{-\infty}^{\infty} n\phi(t)\left[1-\Phi(t)\right]^{n-1}\left[1-\Phi\left(\frac{z_K - \frac{s_m+s_f}{2} - \frac{w_m+w_f}{2} - tr_{\text{ps}}/\sqrt{2}}{\sqrt{1-h^2/2-r_{\text{ps}}^2/2}}\right)\right]dt \\ &= \int_{-\infty}^{\infty} n\phi(t)\left[1-\Phi(t)\right]^{n-1}\left[1-\Phi\left(\frac{z_K - \frac{g_m+g_f}{2} - tr_{\text{ps}}/\sqrt{2}}{\sqrt{1-h^2/2-r_{\text{ps}}^2/2}}\right)\right]dt. \end{aligned} \tag{48}$$

Note that the final result depends only on $g_m$ and $g_f$. Thus, Eq. (48) can be integrated over $g_m$ and $g_f$ (according to their posterior distribution given the family disease history; see Section 6.6) to provide the disease risk probability.

#### 6.4 Excluding high-risk embryos

Here, the density of the score of the selected embryo is given by Eq. (28), which continues to hold, with $c = (s_m + s_f)/2$.

$$f(x_s) = \begin{cases} \dfrac{\frac{1}{r_{\text{ps}}/\sqrt{2}}\phi\left(\frac{x_s}{r_{\text{ps}}/\sqrt{2}}\right)}{1-\Phi\left(\frac{z_q r_{\text{ps}}-c}{r_{\text{ps}}/\sqrt{2}}\right)}\left[1-\Phi\left(\frac{z_q r_{\text{ps}}-c}{r_{\text{ps}}/\sqrt{2}}\right)\right]^n & \text{for } x_s > z_q r_{\text{ps}} - c \\ \dfrac{\frac{1}{r_{\text{ps}}/\sqrt{2}}\phi\left(\frac{x_s}{r_{\text{ps}}/\sqrt{2}}\right)}{\Phi\left(\frac{z_q r_{\text{ps}}-c}{r_{\text{ps}}/\sqrt{2}}\right)}\left\{1-\left[1-\Phi\left(\frac{z_q r_{\text{ps}}-c}{r_{\text{ps}}/\sqrt{2}}\right)\right]^n\right\} & \text{for } x_s < z_q r_{\text{ps}} - c \end{cases} \tag{49}$$

Integrating over all $x_s$, following similar steps as in Section 4, we obtain, denoting by $D_s$ the event that the selected embryo is affected,

$$P\left(D_s \,|\, s_m, w_m, s_f, w_f\right) = \int_{-\infty}^{\infty} \eta(t,\gamma)\xi(t)dt, \tag{50}$$

where we defined

$$\begin{aligned}
\xi(t) &= \phi(t)\left[1-\Phi\left(\frac{z_K - tr_{\text{ps}}/\sqrt{2} - \frac{s_m+s_f}{2} - \frac{w_m+w_f}{2}}{\sqrt{1-h^2/2-r_{\text{ps}}^2/2}}\right)\right] \\
&= \phi(t)\left[1-\Phi\left(\frac{z_K - tr_{\text{ps}}/\sqrt{2} - \frac{g_m+g_f}{2}}{\sqrt{1-h^2/2-r_{\text{ps}}^2/2}}\right)\right], \\
\eta(t,\gamma) &= \begin{cases} \frac{1-[1-\Phi(\gamma)]^n}{\Phi(\gamma)} & \text{for } t<\gamma, \\ [1-\Phi(\gamma)]^{n-1} & \text{for } t>\gamma \end{cases}, \text{ and} \\
\gamma &= \sqrt{2}z_q - \frac{c}{r_{\text{ps}}/\sqrt{2}}.
\end{aligned} \tag{51}$$

Here, Eq. (50) depends on $c, g_m, g_f$, and they must be integrated over to obtain the final disease probability.

#### 6.5 The baseline risk

To compute the relative risk reduction, we need the baseline risk, i.e., the risk when selecting a embryo at random given the parental genetic components. We have

$$\begin{aligned}
P\left(D_s \mid s_m, w_m, s_f, w_f\right) &= P(y_i > z_K) \\
&= P\left(\frac{s_m+s_f}{2} + x_i + \frac{w_m+w_f}{2} + v_i + \epsilon_i > z_K\right) \\
&= P\left(x_i + v_i + \epsilon_i > z_K - \frac{g_m+g_f}{2}\right) \\
&= 1-\Phi\left(\frac{z_K - \frac{g_m+g_f}{2}}{\sqrt{1-h^2/2}}\right).
\end{aligned} \tag{52}$$

The last line holds because $\text{Var}\left(x_i+v_i+\epsilon_i\right) = r_{\text{ps}}^2/2 + (h^2 - r_{\text{ps}}^2)/2 + (1-h^2) = 1-h^2/2$.

#### 6.6 The disease risk conditional on the parental disease status

In subsections 6.3, 6.4, and 6.5, we computed the disease probability under the various strategies given the parental genetic components. For the baseline risk and for the *lowest-risk prioritization* strategy, the risk depended only on $g_m$ and $g_f$. For the *high-risk exclusion* strategy, the risk also depended on $c$. In this section, we compute the posterior probability of these genetic components conditional on the disease status of the parents.

Denote by $D_m$ the indicator variable that the mother is affected (i.e., $D_m = 1$ if the mother is affected and $D_m = 0$ otherwise), and define $D_f$ similarly. The risk of the selected embryo conditional on the parental disease status can be written as

$$
\begin{aligned}
P\left(D_s \mid D_m, D_f\right) &= \iiint dg_m dg_f d_c P\left(D_s \mid g_m, g_f, c, D_m, D_f\right) f(g_m, g_f, c \mid D_m, D_f) \\
&= \iiint dg_m dg_f dc P\left(D_s \mid g_m, g_f, c\right) f(c \mid g_m, g_f) f(g_m, g_f \mid D_m, D_f).
\end{aligned} \tag{53}
$$

The second line of Eq. (53) consists of three terms. The first is $P\left(D_s \mid g_m, g_f, c\right)$, which was computed in the previous subsections for the various selection strategies. Note that we assumed $P\left(D_s \mid g_m, g_f, c, D_m, D_f\right) = P\left(D_s \mid g_m, g_f, c\right)$. This holds because given the genetic components of the parents, their disease status does not provide additional information on the disease status of the children, at least under a model where the environment is not shared (see Section 10). The second term is the density of $c$, which can be similarly written as $f(c \mid g_m, g_f, D_m, D_f) = f(c \mid g_m, g_f)$. The third term is the posterior distribution of $g_m$ and $g_f$ given the parental disease status, $f(g_m, g_f \mid D_m, D_f)$. In the following, we derive the third term and then the second term.

Note that if $P\left(D_s \mid g_m, g_f, c\right) = P\left(D_s \mid g_m, g_f\right)$, as in the case of the baseline risk (Eq. 52) and the *lowest-risk prioritization* (Eq. (48)), the risk of the selected embryo can be simplified by integrating over $c$,

$$
P\left(D_s \mid D_m, D_f\right) = \iint dg_m dg_f P\left(D_s \mid g_m, g_f\right) f(g_m, g_f \mid D_m, D_f). \tag{54}
$$

#### 6.7 The distribution of the parental genetic components given the parental disease status

First, we assume (given that we did not model assortative mating) that given one parent's disease status, his/her genetic component is independent of the spouse's disease status or genetic factors. Thus, the posterior distribution can be factored into

$$
f(g_m, g_f \mid D_m, D_f) = f(g_m \mid D_m) f(g_f \mid D_f). \tag{55}
$$

Next, without loss of generality, we focus on just the mother. To derive the posterior distribution $f(g_m \mid D_m)$ we first need the prior, $g_m \sim N(0, h^2)$.

$$
f_{pr}(g_m) = \frac{1}{h}\phi\left(\frac{g_m}{h}\right). \tag{56}
$$

Next, the likelihood that the mother is affected is

$$
\begin{aligned}
P(D_m = 1 \mid g_m) &= P(y > z_K) \\
&= P(g_m + \epsilon > z_K) \\
&= P(\epsilon > z_K - g_m) \\
&= 1 - \Phi\left(\frac{z_K - g_m}{\sqrt{1-h^2}}\right).
\end{aligned} \tag{57}
$$

Similarly,

$$P(D_m = 0 \,|\, g_m) = \Phi\left(\frac{z_K - g_m}{\sqrt{1-h^2}}\right). \tag{58}$$

Using Bayes' theorem,

$$\begin{aligned} f(g_m \,|\, D_m = 1) &= \frac{P(D_m = 1 \,|\, g_m) f_{pr}(g_m)}{P(D_m = 1)} \\ &= \frac{\left[1 - \Phi\left(\frac{z_K - g_m}{\sqrt{1-h^2}}\right)\right] \frac{1}{h}\phi\left(\frac{g_m}{h}\right)}{K}. \end{aligned} \tag{59}$$

Similarly,

$$\begin{aligned} f(g_m \,|\, D_m = 0) &= \frac{P(D_m = 0 \,|\, g_m) f_{pr}(g_m)}{P(D_m = 0)} \\ &= \frac{\Phi\left(\frac{z_K - g_m}{\sqrt{1-h^2}}\right) \frac{1}{h}\phi\left(\frac{g_m}{h}\right)}{1-K}. \end{aligned} \tag{60}$$

The same results hold for $f(g_f \,|\, D_f = 1)$ and $f(g_f \,|\, D_f = 1)$. We have thus specified the posterior distribution $f(g_m, g_f \,|\, D_m, D_f)$.

#### 6.8 The distribution of the parental mean score given the parental genetic components

The final missing term is $f(c \,|\, g_m, g_f)$. To compute this distribution, we note that $c$, $g_m$, and $g_f$ have a multivariate normal distribution,

$$(c, g_m, g_f) \sim \text{MVN}\left(\begin{pmatrix} 0 \\ 0 \\ 0 \end{pmatrix}, \begin{pmatrix} \frac{r_{\text{ps}}^2}{2} & \frac{r_{\text{ps}}^2}{2} & \frac{r_{\text{ps}}^2}{2} \\ \frac{r_{\text{ps}}^2}{2} & h^2 & 0 \\ \frac{r_{\text{ps}}^2}{2} & 0 & h^2 \end{pmatrix}\right). \tag{61}$$

To explain the above equation, recall that $\text{Var}(c) = r_{\text{ps}}^2/2$ and $\text{Var}(g_m) = \text{Var}(g_f) = h^2$. Then,

$$\begin{aligned} \text{Cov}(c, g_m) &= \text{Cov}\left(\frac{s_m + s_f}{2}, g_m\right) = \frac{1}{2}\text{Cov}(s_m, g_m) \\ &= \frac{1}{2}\text{Cov}(s_m, s_m + w_m) = \frac{1}{2}\text{Var}(s_m) = \frac{r_{\text{ps}}^2}{2}. \end{aligned} \tag{62}$$

A similar result holds for the paternal genetic component. To compute the density of $c$ given $g_m$ and $g_f$, we use standard theory for multivariate normal variables (as in Section 2.1). We have

$$c \,|\, g_m, g_f \sim N(\mu, \sigma^2), \tag{63}$$

with

$$\mu = \frac{r_{\text{ps}}^2}{h^2}\left(\frac{g_m + g_f}{2}\right),$$

$$\sigma^2 = \frac{r_{\text{ps}}^2}{2h^2}(h^2 - r_{\text{ps}}^2). \tag{64}$$

We have thus specified $f(c \,|\, g_m, g_f)$.

##### 6.9 Summary of the computation

In summary, for the *high-risk exclusion* strategy, the probability of disease of the selected embyro given the parental disease status is given by Eq. (53), with $P\left(D_s \,|\, g_m, g_f, c\right)$ given in Eq. (50) and $f(c \,|\, g_m, g_f)$ in Eq. (63). The conditional probability of disease for the *lowest-risk prioritization* strategy and for random selection (the baseline risk) is given by Eq. (54), with $P\left(D_s \,|\, g_m, g_f\right)$ given in Eqs. (48) and Eq. (52), respectively. For all selection strategies, $f(g_m, g_f \,|\, D_m, D_f)$ is given by Eqs. (55), (59), and (60), depending on the particular family history.

Numerically, computing the baseline disease risk requires two integrals (over $g_m$ and $g_f$). Computing the risk for the *lowest-risk prioritization* strategy requires three integrals (over $g_m$, $g_f$, and $t$). Computing the risk for the *high-risk exclusion* strategy requires four integrals (over $g_m$, $g_f$, $c$, and $t$).

#### 7 Two diseases

Prioritizing embryos based on low risk for a target disease may increase risk for a second disease, if that disease is genetically anti-correlated with the target disease. In this section, we develop a model for the PRSs of two diseases in order to investigate this risk.

We denote the variance explained by the scores of the two diseases as $r_1^2$ and $r_2^2$, where disease 1 is the target disease (i.e., embryos are prioritized based on their risk for that disease), and disease 2 is the correlated disease. Denote the genetic correlation between the diseases as $\rho$ (where $\rho < 0$ is the case raising the concern about increasing the risk of the correlated disease), the scores of a child as $s^{(1)}$ and $s^{(2)}$, the scores of the mother as $s_m^{(1)}$ and $s_m^{(2)}$, and the scores of the father as $s_f^{(1)}$ and $s_f^{(2)}$. The vector $(s^{(1)}, s^{(2)}, s_m^{(1)}, s_m^{(2)}, s_f^{(1)}, s_f^{(2)})$ has a multivariate normal distribution, with zero means, and with the following covariance matrix (extending Eq. (9)).

$$\boldsymbol{\Sigma} = \begin{matrix} & s^{(1)} & s^{(2)} & s_m^{(1)} & s_m^{(2)} & s_f^{(1)} & s_f^{(2)} \\ s^{(1)} & r_1^2 & \rho r_1 r_2 & \frac{r_1^2}{2} & \frac{\rho r_1 r_2}{2} & \frac{r_1^2}{2} & \frac{\rho r_1 r_2}{2} \\ s^{(2)} & \rho r_1 r_2 & r_2^2 & \frac{\rho r_1 r_2}{2} & \frac{r_2^2}{2} & \frac{\rho r_1 r_2}{2} & \frac{r_2^2}{2} \\ s_m^{(1)} & \frac{r_1^2}{2} & \frac{\rho r_1 r_2}{2} & r_1^2 & \rho r_1 r_2 & 0 & 0 \\ s_m^{(2)} & \frac{\rho r_1 r_2}{2} & \frac{r_2^2}{2} & \rho r_1 r_2 & r_2^2 & 0 & 0 \\ s_f^{(1)} & \frac{r_1^2}{2} & \frac{\rho r_1 r_2}{2} & 0 & 0 & r_1^2 & \rho r_1 r_2 \\ s_f^{(2)} & \frac{\rho r_1 r_2}{2} & \frac{r_2^2}{2} & 0 & 0 & \rho r_1 r_2 & r_2^2 \end{matrix} \tag{65}$$

In the above covariance matrix, we assumed that the correlation between the scores of the two diseases is also $\rho$. The covariance between parent-child scores for different diseases is half the covariance of the scores within an individual (e.g., see (Karavani et al., 2019)).

Next, we need the density of $\left(s^{(1)}, s^{(2)}\right)$, conditional on $\left(s_m^{(1)}, s_m^{(2)}, s_f^{(1)}, s_f^{(2)}\right)$. We follow a similar procedure as in Section 2.1, and obtain the conditional density of $\left(s^{(1)}, s^{(2)}\right)$ as $\mathrm{MVN}(\boldsymbol{\mu_c}, \boldsymbol{\Sigma_c})$ with

$$\begin{aligned} \boldsymbol{\mu_c} &= \begin{pmatrix} \frac{s_m^{(1)}+s_f^{(1)}}{2} \\ \frac{s_m^{(2)}+s_f^{(2)}}{2} \end{pmatrix}, \\ \boldsymbol{\Sigma_c} &= \begin{pmatrix} \frac{r_1^2}{2} & \frac{\rho r_1 r_2}{2} \\ \frac{\rho r_1 r_2}{2} & \frac{r_2^2}{2} \end{pmatrix}. \end{aligned} \tag{66}$$

We would like to compute the expected increase in risk to become affected by the second disease, given any selection strategy of embryos based on a PRS for the first disease. Solving this problem analytically is beyond the scope of this work. However, the above results imply a method we could use for simulations.

Let us first consider how to draw the average parental scores, which we denote $c^{(1)} = \left(s_m^{(1)} + s_f^{(1)}\right)/2$ and $c^{(2)} = \left(s_m^{(2)} + s_f^{(2)}\right)/2$. The vector $\left(c^{(1)}, c^{(2)}\right)$ has a multivariate normal distribution with zero means (as each parental score has zero mean in the population), and the following covariance matrix. The variances are $\mathrm{Var}\left(c^{(1)}\right) = r_1^2/2$ and $\mathrm{Var}\left(c^{(2)}\right) = r_2^2/2$. The covariance is

$$\begin{aligned} \mathrm{Cov}\left(c^{(1)}, c^{(2)}\right) &= \mathrm{Cov}\left(\frac{s_m^{(1)} + s_f^{(1)}}{2}, \frac{s_m^{(2)} + s_f^{(2)}}{2}\right) \\ &= \frac{1}{2}\mathrm{Cov}\left(s_m^{(1)}, s_m^{(2)}\right) = \frac{\rho r_1 r_2}{2}. \end{aligned} \tag{67}$$

Thus, the covariance matrix is equal to $\boldsymbol{\Sigma_c}$ from Eq. (66) above. This suggests the following simple algorithm for generating the risk scores of the embryos.

$$\begin{pmatrix} c^{(1)} \\ c^{(2)} \end{pmatrix} \sim \text{MVN}\left(\begin{pmatrix} 0 \\ 0 \end{pmatrix}, \begin{pmatrix} \frac{r_1^2}{2} & \frac{\rho r_1 r_2}{2} \\ \frac{\rho r_1 r_2}{2} & \frac{r_2^2}{2} \end{pmatrix}\right),$$
$$\begin{pmatrix} s_i^{(1)} \\ s_i^{(2)} \end{pmatrix} \sim \text{MVN}\left(\begin{pmatrix} c^{(1)} \\ c^{(2)} \end{pmatrix}, \begin{pmatrix} \frac{r_1^2}{2} & \frac{\rho r_1 r_2}{2} \\ \frac{\rho r_1 r_2}{2} & \frac{r_2^2}{2} \end{pmatrix}\right), \quad i = 1, \ldots, n. \tag{68}$$

In our simulations, we select an embryo based on its score for disease 1, according to a selection strategy. We draw the non-score component for disease 1 of the selected embryo as $e^{(1)} \sim N(0, 1 - r_1^2)$, and the liability of the embryo for that disease is then $y^{(1)} = s^{(1)} + e^{(1)}$. We draw the liability for disease 2 of the selected embryo similarly. In our simulations, we draw $e^{(1)}$ and $e^{(2)}$ independently, even though they are correlated. [Any genetic component not included in the score should be correlated between the diseases, and the environments affecting the two diseases may be correlated as well.] We do this because we are only interested in the marginal outcome for each disease separately (i.e., not in the joint outcome of the two diseases). The selected embryo is designated as affected by each disease if the liability of that disease exceeds its respective threshold.

We note that the above model represents the following approximation. As the scores are (noisily) estimating the total genetic effects, the score of one disease is correlated with the non-score genetic component of the other disease. Thus, a more accurate expression for the liability to disease 2 would take into account not only $s_i^{(2)}$ but also $s_i^{(1)}$. However, the dependence is expected to be weak.

### 8 Comparison to previous work

In the "gwern" blog (`https://www.gwern.net/Embryo-selection`), the utility of embryo selection for traits and/or diseases was investigated. For disease risk, a model similar to ours was studied, based on the liability threshold model. However, the model assumed that given the polygenic score, the distribution of the remaining contribution to the liability has unit variance, instead of $1 - r_{\text{ps}}^2$ (the function `liabilityThresholdValue` therein). Further, the blog provided only numerical results, did not consider the *high-risk exclusion* strategy, did not consider the risk reduction conditional on the parental scores or disease status, and did not consider the *per-couple* relative risk reduction. Treff et al. (Treff et al., 2019a) also employed the liability threshold model to evaluate embryo selection for disease risk. However, they did not consider the *high-risk exclusion* strategy, and did not compute analytically the risk reduction. They only provided simulation results for the case when a parent is affected based on an approximate model.

### 9 Simulations

Our analytical results in the above sections provide exact expressions for the relative risk reduction under various settings in the form of integrals, which we then solve numerically. To validate our analytical derivations and the numerical solutions, we also simulated the scores of embryos under each setting, and verified that the empirical risk reductions agree with the analytical predictions.

To simulate the scores of embryos, we used the representation $s_i = x_i + c$, where $(x_1, \ldots, x_n)$ are independent normals with zero means and variance $r_{\text{ps}}^2/2$, and $c \sim N(0, r_{\text{ps}}^2/2)$ is shared across all embryos. Thus, for each "couple", we first draw $c$, then draw $n$ independent normals $(x_1, \ldots, x_n)$, and then compute the score of embryo $i$ as $s_i = x_i + c$, for $i = 1, \ldots, n$. The score of the selected embryo was the lowest among the $n$ embryos in the *lowest-risk prioritization* strategy. For the *high-risk exclusion* strategy, we selected the first embryo with score $s < z_q r_{\text{ps}}$. If no such embryo existed, we selected the first embryo (except for one analysis, in which, if all embryos were high-risk, we selected the embryo with the lowest score.) We then drew the residual of the liability as $e \sim N(0, 1 - r_{\text{ps}}^2)$, and computed the liability as $s^* + e$, where $s^*$ is the score of the selected embryo. If the liability exceeded the threshold $z_K$, we designated the embryo as affected. We repeated over $10^6$ couples, and computed the probability of disease as the fraction of couples in which the selected embryo was affected. We computed the relative risk reduction using Eq. (33).

For the setting when the parental risk scores are given, we computed $c$ as $c = (s_m + s_f)/2$. We specified the maternal score as a percentile $p_m$, such that the score itself was $s_m = z_{p_m} r_{\text{ps}}$, where $z_{p_m}$ is the $p_m$ percentile of the standard normal distribution. We similarly specified the paternal score. The remaining calculations were as above. For the baseline risk, we used the same data, assuming that the first embryo in each family was selected.

When conditioning on the parental disease status, we first drew the three independent parental components, all as normal variables with zero mean. We drew $s_m$ and $s_f$ with variance $r_{\text{ps}}^2$; $w_m$ and $w_f$ with variance $h^2 - r_{\text{ps}}^2$; and $\epsilon_m$ and $\epsilon_f$ with variance $1 - h^2$. We computed the maternal liability as $y_m = s_m + w_m + \epsilon_m$, and designated the mother as affected if $y_m > z_K$. We similarly designated the paternal disease status. We then drew the score of each embyro as $s_i = c + x_i$, where $c = (s_m + s_f)/2$ (using the already drawn parental scores) and $x_i \sim N(0, r_{\text{ps}}^2/2)$, for $i = 1, \ldots, n$, are independent across embryos. We selected one embryo based on the selection strategy, as described above. If $s^*$ is the score of the selected embryo, we computed the liability of the selected embyro as $s^* + (w_m + w_f)/2 + v + \epsilon$, where $v \sim N(0, (h^2 - r_{\text{ps}}^2)/2)$ and $\epsilon \sim N(1 - h^2)$. We designated the embryo as affected if its liability exceeded $z_K$. We tallied the proportion of affected embryos separately for each number of affected parents (0,1, or 2). To compute the baseline risk, we again used the first embryo in each family.

For two diseases, we do not have an analytical solution for the change in risk of the second disease. We thus evaluated the risk using simulations only. We considered the *lowest-risk prioritization* strategy and the case of random parents. For each couple and for each embryo, we generated polygenic scores for the two diseases as outlined in Section 7. We selected the embryo with the lowest score for the target disease, but then considered the score of that embryo for the second, correlated disease. Denote by $s^{*(2)}$ the score of the selected embryo for the second disease. We drew the residual of the liability for the second disease as $e^{(2)} \sim N(0, 1 - r_2^2)$, and the liability of the embryo for that disease was then $s^{*(2)} + e^{(2)}$. If the liability exceeded the threshold of that disease, we designated the embryo as affected. We also repeated for a random selection of an embryo for each couple. We computed the relative risk increase based on the ratio between the risks with or without PRS-based selection.

### 10 Limitations of the model

Our model has a number of limitations. First, our results rely on several modeling assumptions. (1) We assumed an infinitesimal genetic architecture for the disease, which will not be appropriate for oligogenic diseases or when screening the embryos for variants of very large effect. We did not assess the robustness of our theoretical results to deviations from normality in the tails of the distributions of the genetic and non-genetic components (although the good agreement with the simulations based on the real genomic data provide some support that the model is reasonable). (2) Assumption (1) implies that the variance of the scores of children are always half the population variance, regardless of the parental PRSs or the disease considered (Eq. (13)). However, as shown in Chen et al. (2020), the variance of the scores in children can vary across families. On the other hand, Chen et al. also showed (Figure 3C therein) that between-family differences decrease when increasing the number of variants included in the PRS; and, as we showed here, the differences seem to be explained mostly by sampling variance. (3) Our model also assumes no assortative mating, which seems reasonable given that for genetic disease risk, correlation between parents is weak (Rawlik et al., 2019), and given that our previous study of traits showed no difference in the results between real and random couples (Karavani et al., 2019). (4) When conditioning on the parental disease status, we assumed independence between the environmental component of the child and either genetic or environmental factors influencing the disease status of the parents. Family-specific environmental factors were shown to be small for complex diseases (Wang et al., 2017). The influence of parental genetic factors on the child's environment is discussed in the next paragraph. Both of these influences, to the extent that they are significant, are expected to reduce the degree of risk reduction.

Second, we assumed that the proportion of variance (on the liability scale) explained by the score is $r_{\text{ps}}^2$, but we did not specify how to estimate it. Typically, $r_{\text{ps}}^2$ is computed and reported by large GWASs based on an evaluation of the score in a test set. However, the variance that will be explained by the score in other cohorts, using other chips, and particularly, in other populations, can be substantially lower (Martin et al., 2019). Relatedly, the variance explained by the score, as estimated in samples of unrelated individuals, is inflated due to population stratification, assortative mating, and indirect parental effects ("genetic nurture") (Kong et al., 2018; Young et al., 2019; Morris et al., 2020; Mostafavi et al., 2020), where the latter refers to trait-modifying environmental effects induced by the parents based on their genotypes. These effects do not contribute to prediction accuracy when comparing polygenic scores between siblings (as when screening IVF embryos), and thus, the variance explained by polygenic scores in this setting can be substantially reduced, in particular for cognitive traits. However, recent empirical work on within-family disease risk prediction showed that the reduction in accuracy is at most modest (Lello et al., 2020), and within-siblings-GWAS yielded similar results to unrelated-GWAS for most physiological traits (Howe et al., 2021).

Third, we implicitly assumed that polygenic scores could be computed with perfect accuracy based on the genotypes of IVF embryos. However, embryos are genotyped based on DNA from a single or very few cells, and whole-genome amplification results in high rates of allele dropout. Further, embryos are often sequenced to low depth. However, we and others have shown that very accurate genotyping of IVF embryos is feasible (Backenroth et al., 2019; Kumar et al., 2015; Natesan et al., 2014; Treff et al., 2019b; Xiong et al., 2019; Yan et al., 2015; Zamani Esteki et al., 2015). Either way, even if sequencing errors do occur, their effect can be readily taken into account. Suppose that $r_0^2$ is the proportion of variance in liability explained by a perfectly genotyped PRS, and that $r_{impute}^2$ is the squared correlation between the true score and the imputed score of an embryo (which can be estimated experimentally). Then $r_{\text{ps}}^2 = r_0^2 \cdot r_{impute}^2$, where $r_{\text{ps}}^2$ is the variance explained by the observed score, i.e., the index used in our models.

Fourth, we did not model the process of IVF and the possible reasons for loss of embryos. Rather, we assumed that $n$ viable embryos are available that would have led to live birth if implanted. The original number of fertilized oocytes would typically be greater than $n$ (see, e.g., the "gwern" blog for more detailed modeling). Similarly, we did not model the age-dependence of the number of embryos; again, we rather assume $n$ viable embryos are available. Finally, we assumed a single embryo transfer. In principle, transfer of, e.g., two embryos is straightforward to simulate: we can select two embryos based on the selection strategy (e.g., under *lowest-risk prioritization*, select the two embryos with the lowest PRSs). Then, if only one of them is born, we can assume that the child is each of the embryos with probability 0.5. We expect the RRR to somewhat decrease under multiple embryo transfer, for both the *lowest-risk prioritization* and *high-risk exclusion* selection strategies. However, an analytical derivation seems difficult.

Fifth, the residual $e$ in Eq. (2) ($y = s + e$) has a complex pattern of correlation between siblings. As noted in Section 2, $e$ has contributions from both genetic and environmental factors. The genetic covariance between siblings is straightforward to model (as in Section 6). However, the proportion of variance in liability explained by shared environment needs to be estimated and can be large (Lakhani et al., 2019). Further, embryos from the same IVF cycle (when only one is actually implanted) would have experienced the same early developmental environment, and are thus expected to share even more environmental factors, similarly to twins. In the current work, the correlation between non-genetic factors across embryos does not enter our derivations. However, care must be taken in any attempt to model the joint phenotypic outcomes of multiple embryos.

Finally, in this work, we modeled various scenarios for the ascertainment of the parents: either randomly, or based on their scores, or based on their disease status. In future work, it will be interesting to model other settings of family history, such as the presence of an affected child. Further, it is likely that parents will attempt to screen the embryos for more than one disease (Treff et al., 2020). In future work, it will be important to model screening for multiple diseases and compute the expected outcomes.

### 11 Code availability

The R code we used to implement all calculations in this paper and generate the figures can be found at `https://github.com/scarmi/embryo_selection`.

To give two examples, below is an R function that computes the relative risk reduction under the *lowest-risk prioritization* strategy for randomly ascertained parents.

```
library(MASS)
risk_reduction_lowest = function(r,K,n)
{
  zk = qnorm(K, lower.tail=F)
  integrand_lowest = function(t)
    return(dnorm(t)*pnorm((zk-t*sqrt(1-r^2/2)) / (r/sqrt(2)), lower.tail=F)^n)
  risk = integrate(integrand_lowest,-Inf,Inf)$value
  return((K-risk)/K)
}
```

The R function below computes the relative risk reduction under the *high-risk exclusion* strategy (for randomly ascertained parents).

```
risk_reduction_exclude = function(r,K,q,n)
{
  zk = qnorm(K, lower.tail=F)
  zq = qnorm(q, lower.tail=F)
  integrand_t = function(t,u)
    return(dnorm(t)*pnorm((zk-r/sqrt(2)*(u+t))/sqrt(1-r^2),lower.tail=F))
  integrand_u = function(us)
  {
    y = numeric(length(us))
    for (i in seq_along(us))
```

```
    {
      u = us[i]
      beta = zq*sqrt(2)-u
      internal_int1 = integrate(integrand_t,-Inf,beta,u)$value
      denom1 = pnorm(beta)
      if (denom1==0) {denom1=1e-300} # Avoid dividing by zero
      numer1 = 1-pnorm(beta,lower.tail=F)^n
      internal_int2 = integrate(integrand_t,beta,Inf,u)$value
      prefactor2 = pnorm(beta,lower.tail=F)^(n-1)
      y[i] = dnorm(u) * (numer1/denom1*internal_int1 + prefactor2*internal_int2)
    }
    return(y)
  }
  risk = integrate(integrand_u,-Inf,Inf)$value
  return((K-risk)/K)
}
```
